# Assessment of a highly multiplexed RNA sequencing platform and comparison to existing high-throughput gene expression profiling techniques

**DOI:** 10.1101/419838

**Authors:** Eric Reed, Elizabeth Moses, Xiaohui Xiao, Gang Liu, Joshua Campbell, Catalina Perdomo, Stefano Monti

## Abstract

The need to reduce per sample cost of RNA-seq profiling for scalable data generation has led to the emergence of highly multiplexed RNA-seq. These technologies utilize barcoding of cDNA sequences in order to combine samples into single sequencing lane to be separated during data processing. In this study, we report the performance of one such technique denoted as sparse full length sequencing (SFL), a ribosomal RNA depletion-based RNA sequencing approach that allows for the simultaneous sequencing of 96 samples and higher. We offer comparisons to well established single-sample techniques, including: full coverage Poly-A capture RNA-seq and microarray, as well as another low-cost highly multiplexed technique known as 3’ digital gene expression (3’DGE). Data was generated for a set of exposure experiments on immortalized human lung epithelial (AALE) cells in a two-by-two study design, in which samples received both genetic and chemical perturbations of known oncogenes/tumor suppressors and lung carcinogens. SFL demonstrated improved performance over 3’DGE in terms of coverage, power to detect differential gene expression, and biological recapitulation of patterns of differential gene expression from in vivo lung cancer mutation signatures.

## Introduction

Since its inception in 2008, RNA sequencing has become the gold-standard for whole-transcriptome high-throughput data generation (Mortazavi et al., 2008). In addition to RNA transcript expression quantification, RNA-seq allows for more advanced analyses including *de novo* transcriptome assembly (Robertson et al., 2010) and characterization of alternative splicing variants (Bryant et al., 2012). Furthermore, RNA-seq is species agnostic, such that the same library preparation technique may be utilized for humans, mouse, rat, kidney bean, etc. These represent clear advantages over hybridization-based microarray platforms in which individual microarray platforms are designed to quantify specific transcripts for a specific species (Wang et al., 2009). However, one persistent drawback of RNA-seq has been its relatively high cost. The use of classic RNA-seq techniques for experimental designs that require profiling of many samples – especially when the marginal information value of each sample is relatively low, such as in medium- and high-throughput screening applications – can thus present a disqualifying cost burden.

Large-scale projects based on transcriptional profiling of chemical exposure experiments include the Toxicogenomics Project-Genomics Assisted Toxicity Evaluation System (Open TG-GATEs) (Igarashi et al., 2015), the DrugMatrix database (Ganter et al., 2006), and the Connectivity Map (CMap) (Subramanian et al., 2017), among others. Both the TG-GATEs and the DrugMatrix projects used microarrays for expression profiling, which was at the time significantly less costly than full coverage RNA-sequencing, yet still requiring multi-million budgets. Alternatively, the CMap project utilizes the Luminex-1000 (L1000) profiling platform, a bead-based analog expression assay which quantifies 1,058 human transcripts, which are used to impute the expression of 11,350 additional transcripts (Subramanian et al., 2017). This technique is among the least expensive expression assays available, but it is restricted to human screens and it directly profiles only a limited panel of genes. Given the flexibility of RNA-sequencing platforms, highly multiplexed techniques represent a viable alternative for generating transcriptional data from exposure screens, as well as from other experiments that require a large sample size. Therefore, evaluation of the technical validity of specific techniques serves to inform research strategies for a variety of biological inquiries.

The need to reduce the per sample cost of RNA-seq has led to the adoption of barcoding technologies, where cDNA sequences from individual samples are tagged and their libraries are combined and multiplex sequenced in a single lane (Wang et al., 2011). More recently, these techniques have been optimized to allow multiplex sequencing of 96 samples per lane or higher (Hou et al., 2015; Shishkin et al., 2015). Here, we report the results of our effort at optimizing and evaluating one such technique denoted as sparse full length (SFL) sequencing (Shishkin et al., 2015), a ribosomal RNA depletion-based RNA sequencing approach that allows for the simultaneous sequencing of 96 samples and higher. We offer comparisons to well established single-sample techniques, including: full coverage Poly-A capture RNA-seq and microarray, as well as another low-cost highly multiplexed technique known as 3’ digital gene expression (3’DGE) (Asmann et al., 2009). Assessments include comparisons of coverage between the three RNA-sequencing techniques, as well as signal-to-noise and biological recapitulation of gene-level differential signals between treatment groups for the same samples profiled across SFL, microarray, and 3’DGE. For this evaluation study, we generated a set of exposure experiments on immortalized human lung epithelial (AALE) cells (Lundberg et al., 2002) in a two-by-two study design, in which samples received both genetic and chemical perturbations of known oncogenes/tumor suppressors and lung carcinogens (Figure 1). The goal of this report is not only to assess the performance of our optimized highly multiplexed technique, but to inform future research in terms of the strengths and pitfalls of available cost-effective high throughput transcriptomic profiling techniques.

**Figure 1:**
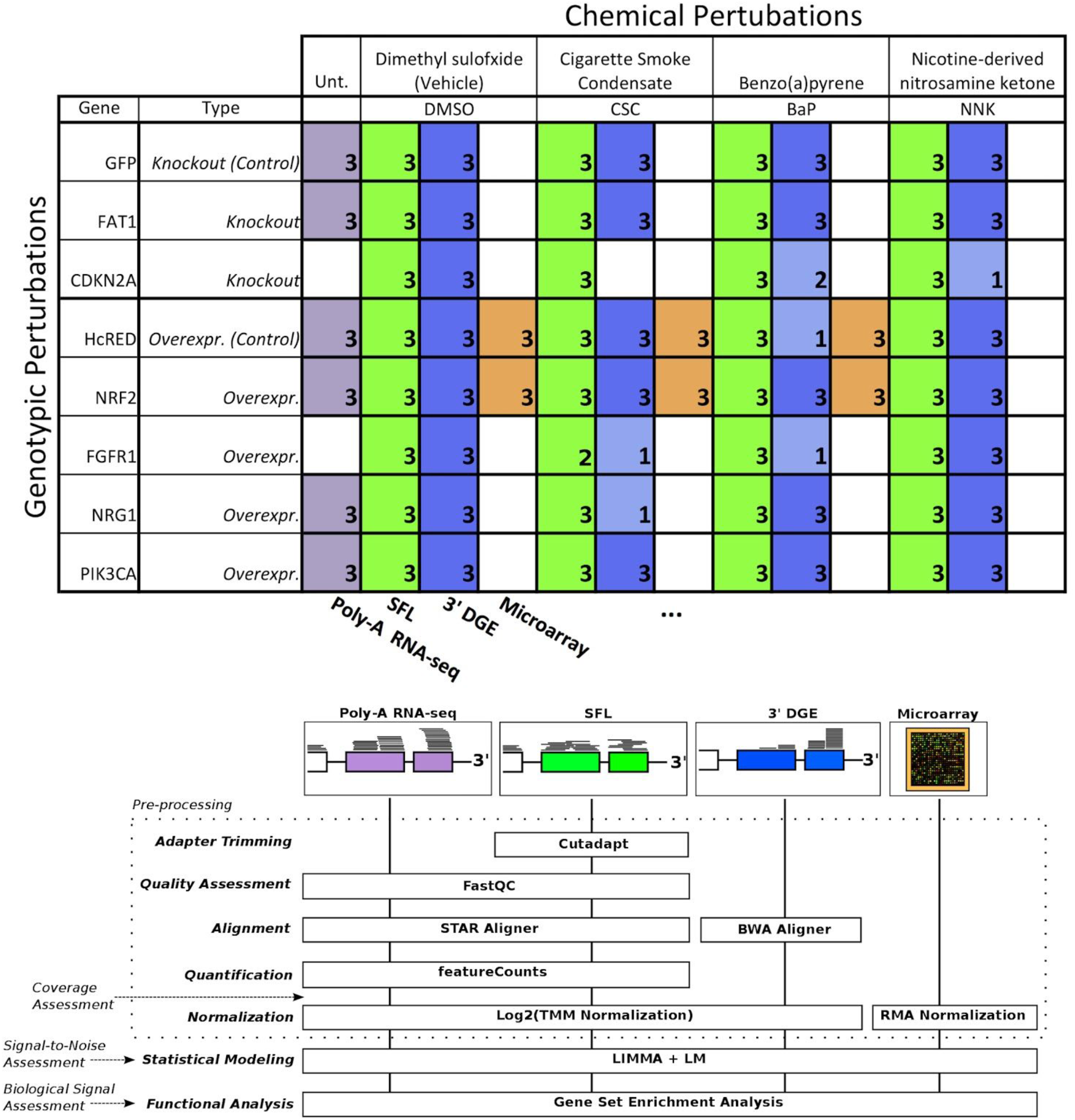
Design of Cross-Platform Experiments and High-throughput Data Processing. Schematic of the number of each pair of genotypic and chemical perturbations, as well as a summary of preprocessing methods used to quantify gene-level expression for each platform. Note that “Unt.” is an abbreviation of “untreated”, denoting that the RNA-seq samples used in this experiment did not receive chemical perturbations. Numbers in each box represent biological replicates of each condition. The color scheme for each platform is consistent throughout this report.

## Materials and Methods

### Samples

Exposure experiments were performed on immortalized human bronchial epithelial cells (AALE). Cells were exposed to both chemical and genotypic perturbations with three replicates per perturbation combination. Cells were thawed from liquid nitrogen and grown up in SAGM small airway epithelial cell growth media (Lonza, Portsmouth NH). Cells were subcultured using Clonetics ReagentPack subculture reagents (Lonza, Portsmouth NH). In preparation for exposure, cells were plated into 24-well plates and allowed to reach confluency for 24 hours. Cell culture media was then replaced, and compounds added at a concentration of 24 μg/ml CSC, 173μM BaP, 490μM NNK or DMSO. NNK and BaP compounds were obtained from Sigma-Aldrich (St. Louis MO) and CSC obtained from Murty Pharmaceuticals (Lexington, KY). Genotypic perturbations included CRISPR knockouts of *FAT1*, and *CDKN2A*, as well as overexpression of *NRF2* (*NFE2L2*), *FGFR1, NRG1* and *PIK3CA*. Cells transfected with a pSpCas9-EGFP (*GFP*) plasmid (PX458) in the absence of sgRNAs were used as controls for the CRISPR perturbations while overexpression of an empty vector containing the reporter HcRed served as control for the overexpression experiments. The same samples were profiled across SFL, microarray, and 3’DGE for a subset of combinations of exposures, though all samples were profiled by SFL. In addition, full coverage poly-A RNA-seq was performed on a separate set of samples for a subset of genotypic exposures, including CRISPR knockouts of *FAT1*, as well as overexpression of *NRF2, NRG1* and *PIK3CA*. These samples did not receive any chemical exposures (Figure 1). Note that in a few cases there was not enough material to perform 3’DGE, as indicated by the sample numbers of certain perturbation combinations.

### Library Preparation

Library preparation for SFL sequencing was carried out based on the published protocol (Shishkin et al., 2015). An edited version of this protocol is available in the Supplementary Methods. RNA was isolated using a standard Qiazol and Qiacube protocol from Qiagen (Valencia, CA). RNA purity was assessed using a NanoDrop spectrophotometer and no samples were excluded from downstream analysis. The dual-barcoded SFL libraries were pooled from 96 individual samples and then sequenced on the Illumina^®^ NextSeq 550 to generate more than 400 million single-end 75-bp reads. Poly-A RNA Sequencing libraries were prepared from total RNA samples using Illumina^®^ TruSeq^®^ RNA Sample Preparation Kit v2 and then sequenced on the Illumina^®^ HiSeq 2500 to generate more than 5 million single-end 50-bp reads per sample. Microarray procedures were performed as described in GeneChip™ WT PLUS Reagent Kit manual and GeneChip™ WT Terminal Labeling and Controls Kit protocol (Thermo Fisher Scientific). The labeled fragmented DNA was generated from 100 ng of total RNA and was hybridized to the GeneChip™ Human Gene 2.0 ST Array. Microarrays were scanned using Affymetrix GeneArray Scanner 3000 7G Plus. 3’DGE library preparation was performed by *Broad Institute, Cambrige, MA, USA*, similar to (Soumillon et al., 2014). Final libraries were purified using AMPure XP beads (Beckman Coulter) according to the manufacturer’s recommended protocol and sequenced on an Illumina NextSeq 500 using paired-end reads of 17bp (read1) + 46bp (read2). Read1 contains the 6-base well barcode along with the 10-base UMI. Across all platforms, the number of samples that were successfully profiled per perturbation combination is shown in Figure 1.

### Data Pre-processing

Affymetrix GeneChip Human Gene 2.0 ST Microarray CEL files were annotated to unique Entrez gene IDs, using a custom CDF file from BrainArray (hugene20st_Hs_ENTREZG_21.0.0) and RMA-normalized. For SFL, adapter sequences were trimmed from raw sequence files using *Cutadapt v1.12*. Quality assessment of trimmed SFL sequence files as well as raw full coverage RNA-seq sequencing files was performed with *FastQC v0.11.5*. Both SFL and RNA-seq reads were aligned to human genome (UCSC RefSeq hg19) with *STAR v2.5.2b* with the non-defulat parameter, *--outSAMtype BAM SortedByCoordinate”* (Dobin et al., 2013). Expression quantification in RefSeq genes was carried out with *featureCounts* (*subread*) *v1.5.0* (Liao et al., 2014). For 3’DGE, pre-quantified gene expression count matrices were obtained from the *Broad Institute, Cambrige, MA, USA*. These reads had been aligned to the transcriptome (UCSC RefSeq hg19), using *BWA aln v0.7.10* with the non-default parameter, *“-l 24”* (Li and Durbin, 2009). Considering that there are 4^10^ (~1.05*10^6^) possible UMIs and the 3’DGE library sizes are on the order of 10^6^ reads, it is highly unlikely for the same UMI to be added to multiple cDNA fragments from the same gene. Therefore, using a custom python program (Soumillon et al., 2014), reads with the same UMI and sample barcode were only counted once per gene. All further data processing and analysis were carried out in *R*.

### Coverage Assessment

Read coverage across the 82 samples, shared between SFL and 3’DGE, as well as all 18 full coverage RNA-seq samples was assessed for library size as well as percentage of the library size that was aligned, uniquely aligned (i.e. reads that only align once in the genome), and counted in the 22,233 genes which were annotated across all three platforms, i.e. the intersection of annotated genes. The full set of counted reads is hereafter referred to as the counted library. Unlike SFL and full coverage RNA-seq, 3’DGE reads are aligned directly to mRNA sequences, such that the reported numbers of counted reads and uniquely aligned reads are the same. To assess the relative distribution of reads across the total set of shared genes, we plotted the cumulative proportion of the sum of reads aligning to individual genes per samples ranked by relative expression across all three platforms. Saturation analysis of the estimated minimum percentage of the counted library size to maximize the number of genes quantified by each platform was performed using a loess fit the gene discovery of 20 subsamplings of the per sample counted libraries. All subsampling analysis was performed using *Subseq v1.8.0*.

Finally, we assessed the relative induction of noise introduced by subsampling progressively larger proportions of the original counted library sizes in each platform, as measured by the principal component error (Heimberg et al., 2016). In order to compare the three platforms assuming equally sized starting library, we repeated the assessment after first subsampling full coverage RNA-seq libraries and 3’DGE libraries to sizes matching that of SFL, the smallest library of the three platforms. This analysis was performed on the 18 samples of like genotypic perturbations, with no chemical treatment in the case of full coverage RNA-seq samples and vehicle DMSO treatment in SFL and 3’DGE samples. Reported values reflect means across 20 iterations of the subsampling and principal component error calculation procedure.

### Signal-to-Noise Assessment

Signal-to-noise was compared among SFL, 3’DGE and microarrays based on four-group ANOVA analysis and two-group differential analysis. In order to estimate signal-to-noise as a means for assessing expected performance when applying standard statistical methods to the data, rather than differential gene expression analysis packages, classic ANOVA was performed for each gene using normalized data across all three platforms, using the *glm* function in R. In this analysis, the signal-to-noise was assessed across like samples undergoing exposure to CSC or DMSO vehicle, as well as genotypic perturbations of *NRF2* overexpression or HcRed control. Thus, the analysis included four independent groups of samples, receiving each combination of chemical (CSC or DMSO) and genotypic (*NRF2* or HcRed) perturbations, with three replicates in each group. Only genes with mean expression ≥ 1 across all 12 samples in both SFL and 3’DGE were included in the analysis (9,813 total genes). Expression levels across SFL and 3’DGE were normalized via trimmed mean of M values (TMM) (Robinson and Oshlack, 2010) scaling and log_2_ counts-per-million transformation. Additionally, two-group differential gene expression analysis was performed for each stratified chemical and genotypic perturbation, using *LIMMA v3.30.7*. That is, differential expression of CSC- vs. DMSO-treated samples, within either HcRed or *NRF2* treatment, as well as differential expression of *NRF2*- vs. HcRed-treated samples, within either DMSO or CSC exposure, was performed. The SFL and 3’DGE count data were transformed for linear modeling based on *voom* (Ritchie et al., 2015). Following modeling, results were restricted to the top 10,000 genes as ranked by median-absolute-deviation (MAD). This heuristic gene filtering procedure was adopted because quantification-based filtering is not applicable to microarray data. This approach follows recommendations detailed in the *LIMMA* manual (Ritchie et al., 2015). All p-values reported from two-group differential analysis are twosided. In both ANOVA and LIMMA analyses, nominal p-values for each gene were corrected for multiple comparisons using the Benjamini-Hochberg procedure (Benjamini and Hochberg, 1995).

### Biological Signal Recapitulation

Two-group differential analysis signatures were compared by pre-ranked gene set enrichment analysis (GSEA) to gene sets derived from published signatures of smoking exposure in the airway from healthy volunteers (Beane et al., 2007; Spira et al., 2004), as well as to gene sets analytically derived from The Cancer Genome Atlas (TCGA) for patients with lung squamous cell carcinoma (LUSC) or lung adenocarcinoma (LUAD). The two smoking gene sets consist of genes reported as either up- or down-regulated in response to smoking in at least one of the two publications, while TCGA gene sets were derived by probing differential expression of individual genes between patients with or without point mutations or copy number alterations (CNA) in genes of interest. These include mutations for the same panel of genes profiled for genotypic perturbations. In addition we include *KEAP1* mutations, a repressor of *NRF2* (Kansanen et al., 2013, 1). Specifically, point mutation signatures were derived from LUSC and LUAD, independently, by performing differential analysis of subjects with and without point mutations in genes of interest, matched for age, sex, and cancer stage. For *NRF2* and *PIK3CA* point mutations were defined at specific mutation hotspots of along the gene body (Figure S2) (Campbell et al., 2016). Likewise, CNA gene signatures were assessed for amplification and deletions of genes of interest by differential analysis, using subjects with zero, one, or two additional copies or deletions of a gene of interest, respectively. All models for mutations and CNA were adjusted for tumor purity, as reported (Campbell et al., 2016). Differential signatures were derived using *LIMMA*. Genes associated with specific mutations or CNA were defined as those with significance and magnitude of the linear model’s genetic alteration coefficient at FDR Q-value < 0.05 and |log2 fold-change| > log_2_(1.5), respectively.

Each of our genotypic perturbation signatures was compared by GSEA to the corresponding TCGA-derived gene sets. For example, the *PIK3CA* overexpression signatures were compared to the gene sets derived from *PIK3CA* mutation and copy number alterations in the TCGA data. To assess the effect of read counts on gene discovery and biological recapitulation of each platform, we compared the differential analysis and GSEA results to that derived from subsampled libraries across full coverage RNA-seq, SFL, and 3’DGE. Similar to coverage assessment, this analysis was performed starting with full libraries across all three platforms, as well as initially subsampling the full coverage RNA-seq and 3’DGE libraries to sizes matching that of SFL. Reported values reflect means from 20 iterations of the subsampling followed by differential analysis and GSEA procedures.

## Results

### Coverage Assessment

Comparison of coverage of the three sequencing platforms, full coverage poly-A RNA-seq, SFL, and 3’DGE, is summarized in Table 1, Figure 2, and Figure S1. Comparison between SFL and 3’DGE included 82 samples each, while full coverage poly-A RNA-seq included all 18 available samples. None of the three platforms demonstrated differences in the library size variability (total number of assigned reads) across samples, although there was a notably high difference between the largest and smallest library size for the SFL samples, with a fold change of 4.3. Fold changes for full coverage RNA-seq and 3’DGE were 1.9 and 2.9, respectively (Table 1, Figure 2A).

**Figure 2:**
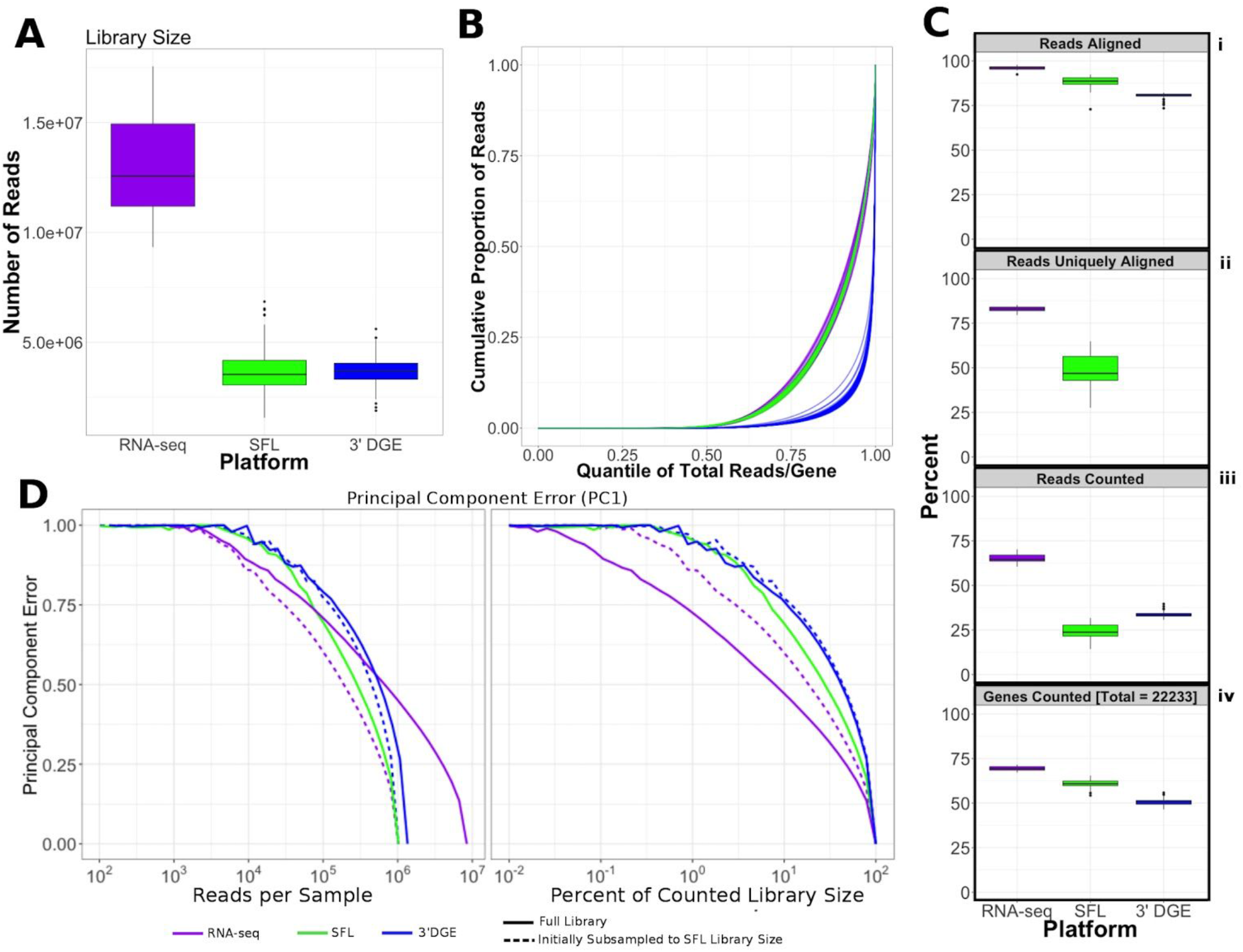
Comparison of Coverage Between Poly-A RNA-seq, SFL, and 3’DGE. A. Boxplots of distribution of library size for each platform.
B. Cumulative distribution of reads assigned to individual genes per sample. The x-axis indicates the quantile for each gene in terms of ranking by relative expression. The y-axis shows the cumulative proportion of total counted reads assigned to these genes, i.e., the running sum of reads divided by the total number of reads across all genes.
C. The top 3 boxplots show the percentage of reads aligned (i), uniquely aligned (ii), and counted(iii) relative to the total library size for each platform. The bottom boxplot (iv) shows the proportion of genes with counts > 1, for protein-coding genes annotated across all 3 platforms (18,488). For Figure 2Cii, “Reads Uniquely Aligned” is not shown for 3’DGE because “Reads Uniquely Aligned” and “Reads Counted” are the same values as a result of the data pre-processing protocol, specific to 3’DGE (see Methods). Counts values for these percentages are given in Figure S1A.
D. Analysis of the principal component error of subsampled counted library sizes for full coverage poly-A RNA-seq, SFL, and 3’DGE for principal component 1. Results for principal component 2-5 is shown in Figure S1D. Initial subsamples of Poly-A RNA-seq and 3’DGE to the SFL library size are also given as dotted lines.

**Table 1:**
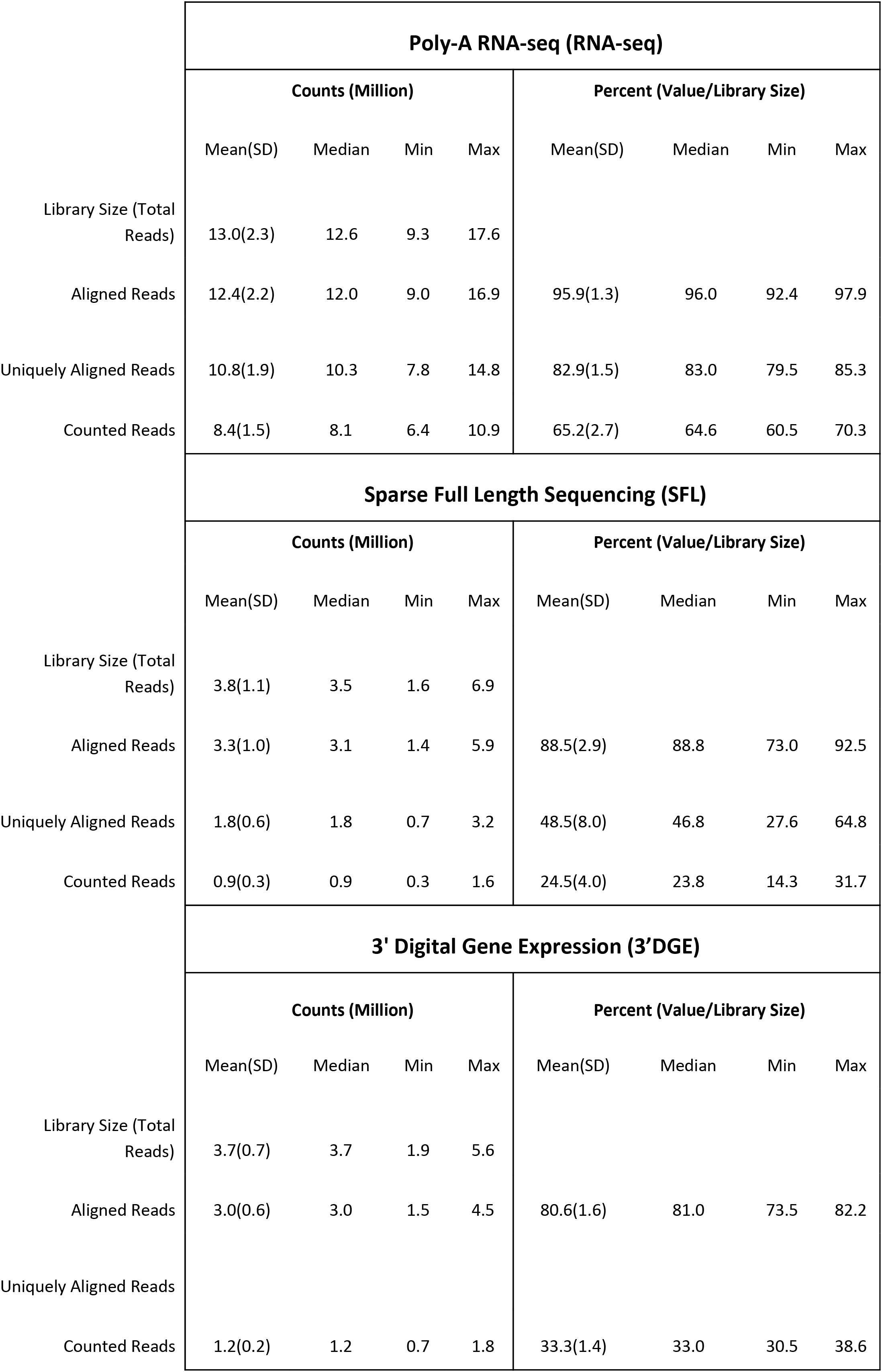
Comparison of Read Assignment Between Full Coverage Poly-A RNA-seq, SFL, and 3’DGE.

| Poly-A RNA-seq (RNA-seq) |  |  |  |  |  |  |  |  |
| --- | --- | --- | --- | --- | --- | --- | --- | --- |
| Counts (Million) |  |  |  |  | Percent (Value/Library Size) |  |  |  |
|  | Mean(SD) | Median | Min | Max | Mean(SD) | Median | Min | Max |
| Library Size (Total Reads) | 13.0(2.3) | 12.6 | 9.3 | 17.6 |  |  |  |  |
| Aligned Reads | 12.4(2.2) | 12.0 | 9.0 | 16.9 | 95.9(1.3) | 96.0 | 92.4 | 97.9 |
| Uniquely Aligned Reads | 10.8(1.9) | 10.3 | 7.8 | 14.8 | 82.9(1.5) | 83.0 | 79.5 | 85.3 |
| Counted Reads | 8.4(1.5) | 8.1 | 6.4 | 10.9 | 65.2(2.7) | 64.6 | 60.5 | 70.3 |
| Sparse Full Length Sequencing (SFL) |  |  |  |  |  |  |  |  |
| Counts (Million) |  |  |  |  | Percent (Value/Library Size) |  |  |  |
|  | Mean(SD) | Median | Min | Max | Mean(SD) | Median | Min | Max |
| Library Size (Total Reads) | 3.8(1.1) | 3.5 | 1.6 | 6.9 |  |  |  |  |
| Aligned Reads | 3.3(1.0) | 3.1 | 1.4 | 5.9 | 88.5(2.9) | 88.8 | 73.0 | 92.5 |
| Uniquely Aligned Reads | 1.8(0.6) | 1.8 | 0.7 | 3.2 | 48.5(8.0) | 46.8 | 27.6 | 64.8 |
| Counted Reads | 0.9(0.3) | 0.9 | 0.3 | 1.6 | 24.5(4.0) | 23.8 | 14.3 | 31.7 |
| 3' Digital Gene Expression (3'DGE) |  |  |  |  |  |  |  |  |
| Counts (Million) |  |  |  |  | Percent (Value/Library Size) |  |  |  |

|  | Mean(SD) | Median | Min | Max | Mean(SD) | Median | Min | Max |
| --- | --- | --- | --- | --- | --- | --- | --- | --- |
| Library Size (Total Reads) | 3.7(0.7) | 3.7 | 1.9 | 5.6 |  |  |  |  |
| Aligned Reads | 3.0(0.6) | 3.0 | 1.5 | 4.5 | 80.6(1.6) | 81.0 | 73.5 | 82.2 |
| Uniquely Aligned Reads |  |  |  |  |  |  |  |  |
| Counted Reads | 1.2(0.2) | 1.2 | 0.7 | 1.8 | 33.3(1.4) | 33.0 | 30.5 | 38.6 |

Unsurprisingly, full coverage poly-A RNA-seq generated the largest library size, while the SFL and 3’DGE libraries were of comparable size (Figure 2A). Furthermore, full coverage poly-A RNA-seq yielded the highest percentage of reads aligned to the genome, followed by SFL and 3’DGE (Table 1, Figure 2Ci, Figure S1A). The lower mapping rate of 3’DGE is most likely due to the lower read quality scores of 3’DGE compared to full coverage RNA-seq and SFL (Figure S1B). The mean percentage of reads with Phred quality scores greater than 20 (Q20) was only ~88% for 3’DGE, compared to ~100% for both full coverage RNA-seq and SFL. The relative 5’-3’ transcript coverage for each sample across all three platforms is shown in Figure S1F. As expected, reads alignments were skewed towards the 3’ end of transcripts for 3’DGE, while we did observe relatively uniform coverage along the transcript for full coverage RNA-seq and SFL.

For SFL there was a clear drop-off when going from percentage of aligned reads to percentage of uniquely aligned reads due to ribosomal RNA (rRNA) contamination of the SFL samples (Figure 2Cii). The majority of reads aligning to ribosomal regions specifically align to RNA28S (Figure S3). For 3’DGE, unique UMIs are aligned directly to transcript sequences and not to the whole genome, such that the number of uniquely aligned reads and reads counted in transcripts are the same (Figure 2Cii-iii) (Morrissy et al., 2009). The percentage of reads that are counted in transcripts is greatest for full coverage poly-A RNA-seq (mean percentage of total library size: 65.2%), followed by 3’DGE (33.3%), and SFL (24.5%). However, while the counted read library size is greater for 3’DGE than for SFL, more genes were quantified by SFL than by 3’DGE (Figure 2Civ) (counts > 0 across all samples for 22,233 genes shared across all three platforms,). A median of 60.9% and 50.5% genes were quantified by SFL and 3’DGE, respectively. The number of genes quantified was near the saturation point for each platform, such that this discrepancy is not due to read depth of each platform (FigureS1C). The reason for the low gene discovery of 3’DGE is further illustrated in Figure 2B, where it is shown that the reads are more evenly distributed across the 22,233 genes by SFL than by 3’DGE, with the cumulative distribution of reads counted in individual genes nearly identical in SFL and full coverage poly-A RNA-seq.

The principal component (PC) error was estimated for each platform for different subsamples of the full counted library size. The first PC is shown in Figure 2D, while the second through the fifth PCs are shown in Figure S1D. We observe that as the counted library size increases, the PC error decreases at the fastest rate for full coverage RNA-seq, followed by SFL, then 3’DGE. Though these differences are more prominent when comparing full coverage RNA-seq to either SFL or 3’DGE, we do observe that when down-sampling from 10% to 100% of the counted library size, the PC error decreases at a consistently faster rate for SFL than for 3’DGE. Initially subsampling full coverage RNA-seq and 3’DGE to match the full SFL counted library size does not change the results. The same trend is also observed in the cumulative variance explained by each successive PC across full coverage RNA-seq, SFL, and 3’DGE (Figure S1E).

In summary, despite lower overall counted library size due to ribosomal RNA contamination, SFL demonstrates greater coverage in low-to-medium expressed genes than 3’DGE, comparable to full coverage poly-A RNA-seq. Consequently, the transcriptional signal captured by the SFL libraries are more robust to subsampling of the data compared to 3’DGE as measured by the principal component error.

### Signal-to-Noise Evaluation

Differential expression models comparing experimental groups of matched samples was performed in SFL, microarray, and 3’DGE and the corresponding signal-to-noise scores were compared pairwise between platforms (Figure 3). Samples shared across the three platforms include three replicates for each of four experimental groups, corresponding to *NRF2* overexpression or HcRed vehicle, as well as CSC chemical exposure or DMSO vehicle (Figure 1). Signal-to-noise was assessed by a four-group comparison with classic ANOVA (Figure 3A-D), as well as by stratified two-group differential analyses using *LIMMA* (Figure 3 E-F).

**Figure 3:**
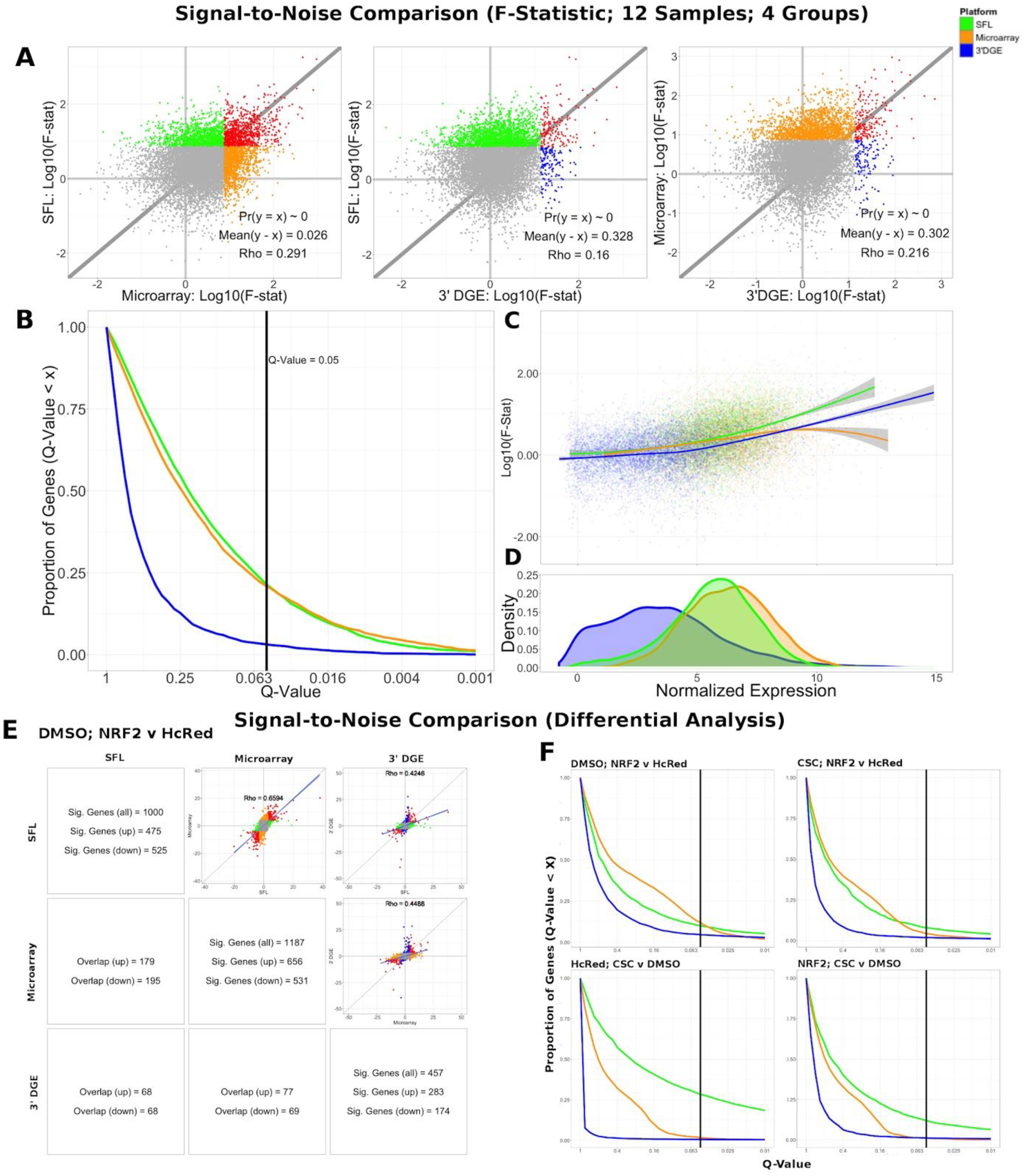
Signal-to-Noise Comparison Between SFL, Microarray, and 3’DGE. A. Scatterplots comparing the log10(F-Statistics) from ANOVA models comparing four n=3 groups (HcRed:DMSO, HcRed:CSC, NRF2:DMSO, and NRF2:CSC). The grey line shows y=x. The platform with the higher mean log10(F-Statistic) is plotted on the y-axis. Also, included are the p-value and difference in mean between each bi-platform comparison from paired t-testing, as well as the squared correlation coefficient. P-values ~ 0 are less than 0.01. Color of indicate genes discovered by individual platforms (green, orange, or blue), neither platform (grey), and both platforms (red).
B. Plot of the Discovery Rate versus FDR Q-Value from threshold for each platform from four group ANOVA models. The x-axis is plotted on a −log10 scale. The vertical line is indicative of a Q-value threshold of 0.05.
C. Loess fit of the log10(F-Statistic) versus median normalized expression from four group ANOVA models.
D. Distribution of mean normalized expression across all three platforms.
E. Comparison of gene discovery (FDR Q-Value < 0.05) by differential analysis with limma, comparing normalized gene expression between DMSO:NRF2 and DMSO:HcRed, including the raw discovery rates, discovered gene overlap, and linear fits, comparing test statistics from each platform. Genes that are discovered by more than 1 platform are shown in red in the scatterplots. Additional comparisons are shown in Figure S5.
F. Plot of the Discovery Rate versus FDR Q-Value from threshold for each platform from two group differential analyses. The x-axis is plotted on a −Log10 scale. The vertical line is indicative of a Q-value threshold of 0.05.

We compared the log_10_ F-statistics between ANOVA models across all three platforms (Figure 3A). Overall, the distribution of F-statistics is most similar between SFL and microarrays, with a Pearson correlation of 0.291. Though statistically significant (p < 0.01), the corresponding mean difference between log_10_ F-statistics is only 0.026. The mean differences of the log_10_ F-statistics between SFL and 3’DGE, and between 3’DGE and microarray are 0.328 and 0.302, respectively, and the corresponding Pearson correlations are 0.160 and 0.216, respectively. These results are consistent with the discovery rates estimated for different FDR Q-value thresholds (Figure 3B). For example, at the FDR Q-value threshold of 0.05, the discovery rates of SFL and microarray are almost identical, 0.214 (2083 genes), 0.209 (2038 genes), respectively, while the discovery rate of 3’DGE is much smaller 0.032 (310 genes).

Loess regression of the log_10_ F-statistics as a function of mean gene expression shows that the statistical signal increases with mean normalized expression. This trend is consistently positive for both SFL and 3’DGE, while leveling off at the most highly expressed genes in microarrays (Figure 3C). Furthermore, SFL signal is greater than 3’DGE signal at all levels of mean expression (Figure 3C). In agreement with the results from coverage comparison, the distribution of mean normalized expressions in 3’DGE is smaller than that of SFL, while SFL is comparable to that of microarray (Figure 3D). Adherence to assumption of normality, assessed through a Shapiro-Wilk test, is also associated with higher mean normalized expression (Figure S4).

The results of the comparisons of the two-group differential analyses across all three platforms were generally congruous with those of the four-group ANOVA analyses (Figure 3E-F, Figure S5, Figure S6). In all four two-group comparisons, the correlation of test statistics is closest between microarray and SFL results, followed by 3’DGE versus microarray results, and 3’DGE versus SFL. For example, in the DMSO-stratified, *NRF2* versus HcRed analysis, estimates of the Pearson correlations of test statistics are 0.66, 0.45, and 0.43, respectively (Figure 3E). The discovery rate of 3’DGE is the lowest across all four differential analyses, while the discovery rate of SFL is higher in three out of four of these analyses (Figure 3F, Figure S5, Figure S6).

In summary SFL demonstrated greater statistical power than 3’DGE to detect differentially expressed genes, and its results more closely matched those in microarrays.

### Biological Signal Recapitulation Evaluation

To evaluate the ability of each platform to recapitulate biologically relevant results, we utilized previously published signatures of smoking exposure in lung (Beane et al., 2007; Spira et al., 2004), as well as differential signatures derived from the TCGA LUSC and LUAD datasets associated with mutations of the genes over-expressed in our experiments. From each of these signatures two gene sets were extracted, one of genes positively associated and one of genes negatively associated to the variable of interest. These gene sets were then tested via pre-ranked gene set enrichment analysis against each of our differential analysis results (CSC *vs*. DMSO, stratified by *NRF2* or HcRed perturbation; *NRF2* vs. HcRed, stratified by CSC or DMSO perturbation). The enrichment results with respect to both the smoking exposure signatures and the TCGA mutations are summarized in Figure 4A, and further detailed in Figure S5, and confirm the highest sensitivity of microarrays, followed by SFL and 3’DGE.

**Figure 4:**
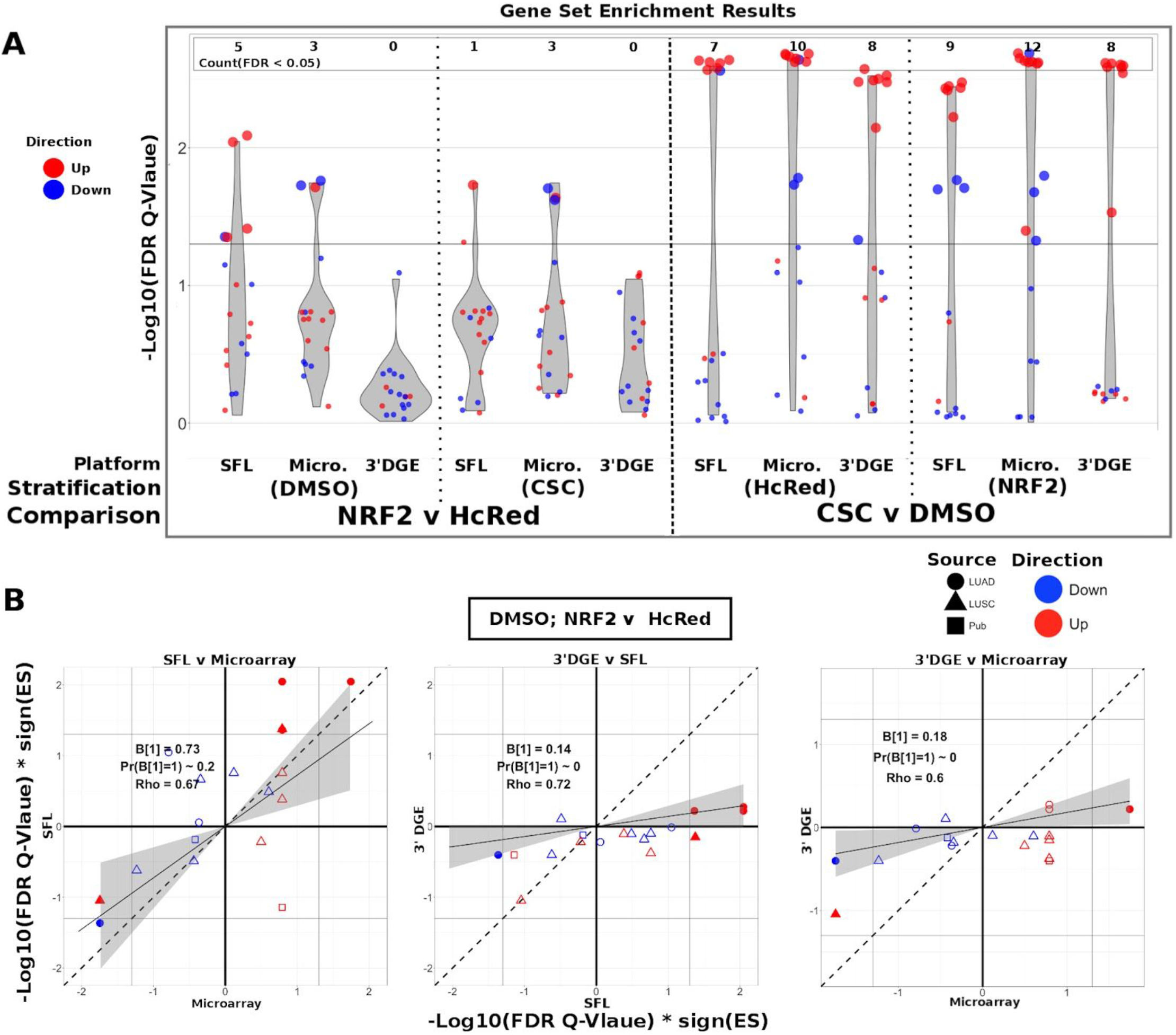
Comparison of Gene-set enrichment of Smoking and Gene Mutation Signatures across SFL, 3’DGE and Microarray. A. Violin plots of the −Log10(FDR Q-Value) from gene set enrichment analysis of TCGA-derived gene-sets with respect genotypic perturbations (left) and chemical perturbations (right) differential signatures across like samples within SFL, Microarray, and 3’DGE. Each column corresponds to differential signatures comparing genotypic or chemical perturbation groups, stratified by a single chemical or genotypic perturbation group, respectively, e.g. the left-most column shows the enrichment results with respect to the “DMSO-treated; NRF2 *vs*. HcRed” signature within the samples (*stratum*) in SFL data. Specific results for TCGA-derived genes sets are shown in Figure S7.
B. Comparison of the gene set enrichment results between SFL, microarray and 3’DGE with respect to the “DMSO-treated; NRF2 *vs*. HcRed” differential signature. Shown are the transformed FDR Q-values of the TCGA-derived gene sets corresponding to mutations of NRF2 and CNA of KEAP1. The |–Log10(FDR Q-Values)| corresponding to the FDR<0.05 significance thresholds are shown as vertical and horizontal gray lines for the y and x-axes, respectively. Points of gene sets whose enrichment meets this threshold in either of the two platforms are filled in. Colors and shape of points denote direction and source of the gene set, respectively. Additional results for chemical and genotypic perturbation signatures are shown if Figure S8.

The set of genes up-regulated in “smokers *vs*. non-smokers” was found to be significantly (FDR Q-value < 0.05) enriched in all “CSC *vs*. DMSO” signatures, within both genotypic stratifications for all three platforms. Conversely, the set of down-regulated genes in “smokers *vs*. non-smokers” was only enriched in the microarray signature of “*NRF2* over-expressed; CSC *vs*. DMSO” (Figure S7).

The enrichment results of TCGA-derived gene sets with respect to differential signatures of genotypic perturbations were in agreement with the gene-level results, in that they consistently demonstrated smaller discovery rates by 3’DGE than by SFL or by microarrays (Figure 4A). For example, the significantly enriched gene sets in “DMSO-treated; *NRF2 vs*. HcRed” differential signatures across all three platforms are highlighted in Figure S7. The number of gene sets enriched in microarray, SFL, and 3’DGE platforms are five, three, and zero, respectively.

In addition to comparing which gene sets were significantly enriched in individual differential signatures, we compared the relative statistical signal of these enrichments. To this end, we transformed the permutation-based FDR Q-values by taking the negative Log_10_ and multiplying by the direction of the enrichment score (ES), −Log10(FDR Q-values)*sign(ES). For each two-platform comparison, we fit a regression model through the origin. Since consistent results across platforms would result in a model fit close to the identity line, *y* = *x*, we tested whether the slope coefficient equaled 1 (i.e. B_1_ = 1). Figure 4B shows these results for each of the three comparisons of the *NRF2* and *KEAP1* mutation-based gene sets enrichment against the “DMSO-treated; *NRF2 vs*. HcRed” signatures. In all three comparisons, microarrays have the highest measured enrichment signal, followed by SFL and 3’DGE, however the difference between microarray and SFL results is not significant, B_1_ = 0.73; p-value = 0.2. The coefficients for both of the comparisons to 3’DGE, are highly skewed in favor of microarray and SFL, B_1_ = 0.18 and 0.14, respectively. Both of these comparisons are highly significant with p-values < 0.01. Comparison of the enrichment results for other differential signatures show similar trends (Figure S8).

Next, we compared enrichment results with respect to all genotypic perturbation signatures between SFL and 3’DGE (Figure 5A; Figure S9A). Each comparison (i.e., each point in the plot) denotes gene set enrichment results with respect to genotypic perturbations within each of the four chemical exposures, DMSO, CSC, BaP, and NNK. Gene sets were tested for enrichment against concordant differential signatures, e.g., the *PIK3CA* mutation-derived gene set was tested against the “*PIK3CA vs*. HcRed” signatures. As in the previous analysis, the permutation-based enrichment FDR Q-values were transformed by −Log10(FDR Q-values)*sign(ES). In the “DMSO-treated; genotypic perturbation *vs*. control” signatures, we observe that the gene set enrichment is generally more significant for SFL than for 3’DGE (B_1_ = 0.63; p-value < 0.01; Figure 5A). The results obtained in CSC- and NNK-treated signatures, demonstrate concordance to these results (B_1_ = 0.65; p-value = 0.03 and B_1_ = 0.60; p-value = 0.01, respectively). The BaP-treated results are less comparable since only one genotypic perturbation signature, “*FAT1 vs*. GFP”, is available for this stratification (Figure S9A).

**Figure 5:**
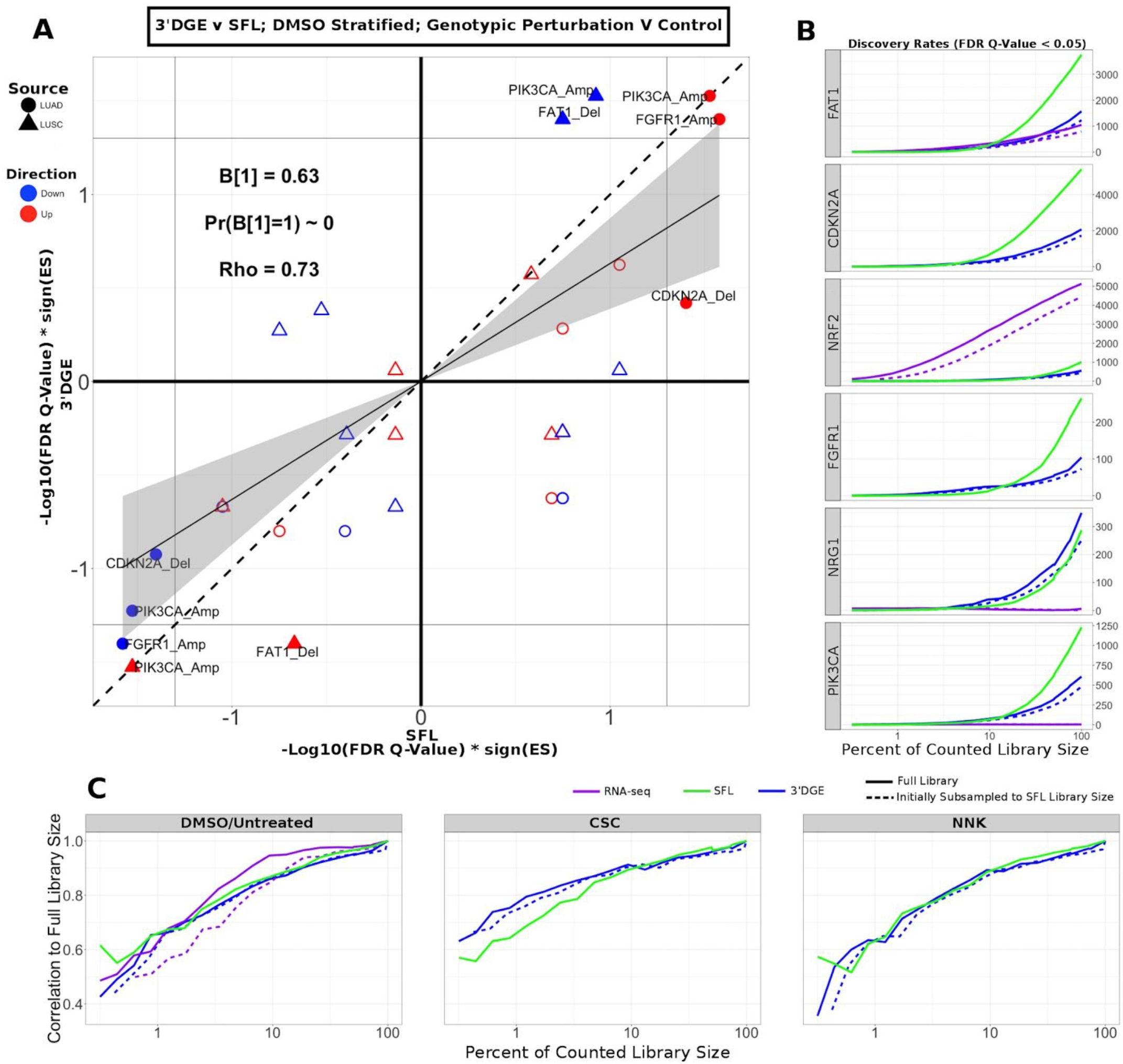
Comparison of Gene-set enrichment of Gene Mutation Signatures across SFL and 3’DGE. A. Comparison of the gene set enrichment results between SFL and 3’DGE with respect to the “DMSO-treated; genotypic perturbation *vs*. control” differential signatures. Points indicate gene set enrichment against concordant signatures, e.g., PIK3CA mutation and CNA gene sets against the “PIK3CA *vs*. HcRed” differential signatures. Shown are the transformed FDR Q-values from permutation-based testing by pre-ranked GSEA. |–Log10(FDR Q-Values)| corresponding to the FDR=0.05 significance thresholds are shown as vertical and horizontal gray lines for the y and x-axes, respectively. The names of the gene sets whose enrichment meets this threshold in either of the two platforms are shown and their points are filled in. Colors and shape of points denote direction and source of the gene set, respectively. Additional results for CSC, NNK, and BaP stratified genotypic perturbation signatures, as well as comparisons between full coverage RNA-seq and either SFL and 3’DGE are shown in Figure S9.
B. Discovery rates for genotypic perturbations across full coverage poly-A RNA-seq, SFL, and 3’DGE, for chemically untreated (full coverage RNA-seq) and DMSO treated (SFL and 3’DGE) samples. Results demonstrate full counted library size, as well as subsampled libraries.
C. Correlation between transformed FDR Q-values from gene set enrichment at different subsamples of each platform and the results from the full counted library size. Shown are the results from genotypic perturbations from untreated (full coverage RNA-seq)/DMSO treated (SFL and 3’DGE), CSC, and NNK chemically treated samples.

Additionally, we compared our differential signatures to available full coverage poly-A RNA-seq genotypic perturbations (Figure S9B), although these results are considered less comparable because of differences in experimental set-up. In particular, in the full coverage poly-A RNA-seq experiments the genotypic perturbations were performed on untreated rather than DMSO-treated cell lines (Figure 1).

The effect on discovery rate by subsampling the data across all three platforms is shown in Figure 5B. Generally, we did not observe a plateauing of discovery rate, where the number of detected genes plateaus near full counted library size. When comparing the correlation between GSEA results on subsampled data we observe similar trends across full coverage RNA-seq, SFL, and 3’DGE (Figure 5C). Initial subsampling of full coverage RNA-seq and 3’DGE to the SFL counted library size did not change the analysis results.

In summary, differential analysis of molecular and genotypic perturbations with SFL recapitulates biologically meaningful signal of gene sets derived from high coverage in vivo data sets. This performance is comparable to both 3’DGE and microarray.

## Discussion

The goal of this study was to evaluate the performance of SFL sequencing, a low-cost method for performing highly multiplexed RNA-seq, and to compare it to other high-throughput gene expression profiling platforms. The development of such methods would be instrumental to the generation of large-scale perturbation screens based on in-vitro models. The reduction of the cost per profile would make it feasible to significantly increase the number of replicates and conditions to be profiled, including multiple time points, concentrations, and biological models, and thus would support a more in-depth investigation of the heterogeneity of the biological response to different exposures. It would also support the development of more accurate predictive models of the adverse or therapeutic outcomes of various exposures. Finally, insights gained from our study will also inform the design of protocols for single cell RNA-sequencing (Eberwine et al., 2014), given their reliance on highly-multiplexed libraries.

In addition to SFL, the platforms included in this analysis were 3’DGE, an alternative highly multiplexed sequencing platform, Affymetrix GeneChip Human Gene 2.0 ST Microarray, an analog expression platform, and full coverage poly-A capture RNA-seq. The cost per sample for SFL and 3’DGE was ~$50, a 10-fold decrease from that of full coverage RNA-seq, $500, and a 7-fold decrease from that of the microarray, $350 USD. Throughout this analysis we demonstrate comparable performances of SFL and 3’DGE to these more expensive platforms. Furthermore, in this analysis we consistently find evidence that SFL outperforms 3’DGE.

Performance was assessed in terms of coverage, signal-to-noise, and recapitulation of expected biological signal derived from independently generated, publicly available data collected from human subjects. Coverage was assessed by comparing the three digital expression platforms, while signal-to-noise and biological recapitulation was assessed by comparing SFL, 3’DGE, and microarrays. Microarray expression quantification has been shown to be highly correlated with qRT-PCR, especially when processed with updated probe set annotations, utilized in this analysis (Sandberg and Larsson, 2007). Chemical and molecular perturbations were carried out in the same samples, and concurrently profiled by SFL, 3’DGE, and microarrays. We also leveraged previously generated full coverage poly-A RNA-seq profiles from similar perturbations of AALE cell lines.

For coverage assessment, performance was evaluated in terms of the distribution of total reads, or library size, that were aligned to the human genome, and further quantified in annotated genes. The best performance was expected in full coverage poly-A RNA-seq, given that this is the most well-established technique and has by far the highest sequencing depth. This was confirmed, as full coverage poly-A RNA-seq was measured to have the highest per sample library size, percentage of aligned reads, percentage of uniquely aligned reads, and percentage of counted reads (Figure 2, Figure S1). The coverage performance of SFL suffered as a result of rRNA contamination, where as many as 53% of the total library size per sample was assigned to ribosomal regions of the genome (Figure S3).

3’DGE is a poly-A capture technique, therefore ribosomal depletion is not a possible pitfall. 3’DGE generates a short nucleotide tags from transposon-based fragmentation, which are enriched for 3’ adjacent sequences of a given transcript (Soumillon et al., 2014). Since many transcripts of the same gene generate identical sequence tags, unique molecular identifiers (UMIs) are used to distinguish between unique reads and duplicate reads generated from PCR amplification. Although mRNA fragment duplication occurs with any RNA-seq protocol, the impact of this artifact on downstream analyses is negligible for techniques, such as SFL, which generate more complex sequence libraries (Parekh et al., 2016).

3’DGE sequences were aligned directly to human mRNAs, rather than the whole genome. Therefore, percentages of reads aligned and reads counted (Figure 2Ci,iii) reflect the percentages of these non-unique UMIs that align to at least one gene and the number of unique UMIs that align to only one gene, respectively. We observe that the percentage of counted reads is greater for 3’DGE than SFL, which is explained by a loss of reads to rRNA contamination in SFL. However, we observe notably more genes quantified by SFL than by 3’DGE (Figure 2B, Figure 2Civ), which indicates that more reads are assigned to fewer genes in 3’DGE compared to SFL, as well as to full coverage RNA-seq (Figure 2C). Although rRNA contamination is a potential drawback of any ribosomal depletion RNA-sequencing technique, the extent of ribosomal contamination is variable, and could be potentially improved by further optimization of the library preparation protocol.

The difference in distribution of reads across shared genes between SFL and 3’DGE likely explains the difference in information retained by subsampling as measured by principal component error. We consistently observe that, as the counted library size increases, the rate of principal component error decreases faster for SFL than 3’DGE (Figure 2D, Figure S1D). This is unsurprising considering that not only are considerably fewer genes quantified by SFL compared to 3’DGE, but there is also no discernable difference between the rate of genes counted as a function of counted library size between the two platforms (Figure S1C). As we subsample the counted libraries, though we may lose the same number of genes between SFL and 3’DGE, the percent of genes lost, and consequently the information lost, will be greater for 3’DGE than SFL. Furthermore, this more even read distribution likely explains the improved performance of SFL over 3’DGE in statistical signal. In particular, our signal-to-noise evaluation shows consistently higher gene-level statistical signal from SFL and microarray experiments than from 3’DGE experiments (Figure 3). These differences appear to be driven by the differences in the relative quantification of genes, given that statistical signal is positively associated with mean gene expression for each platform, and 3’DGE experiments showed lower gene-level quantification than SFL and microarrays (Figure 3C-D). We observe similar cross-platform relationships in the two-group differential analyses (Figure 3E-F).

The gene set-based enrichment results are consistent with those from signal-to-noise analyses. In every comparison of enrichment scores between SFL and 3’DGE, we observe generally higher gene set enrichment with respect to the SFL-derived signatures (Figure 4, Figure 5A, Figure S8, Figure S9). The gene sets were selected to represent known biological responses to the profiled perturbations, and thus their enrichment with respect to the perturbation signatures are expected to be true positives.

The enrichment results confirm this expectation. For example, in the signatures of *NRF2* overexpression, we consistently observe enrichment of the gene sets derived from *NRF2* amplifications and *KEAP1* deletions, each of which should increase *NRF2* activity (Figure S7) (Kansanen et al., 2013). Similarly, we observe significant concordant enrichment of the gene sets derived from *NRF2* and *KEAP1*-dysregulated lung tumors in the signature of CSC exposure, suggesting that the *NRF2* pathway is activated by CSC exposure in vitro (Figure S7), which has been previously reported (Adair-Kirk et al., 2008). Interestingly, these results demonstrate that the activation of the *NRF2* pathway in normal airway epithelial cells *in vitro* (by ectopic expression of the gene or by CSC treatment) is concordant with the activation of *NRF2* by somatic genome alterations in lung tumors, a finding that, to the best of our knowledge, has not been previously observed.

Possible sources of technical variability in this study are the different sequencing platforms, service providers, and read lengths. However, when subsampling the 3’DGE and SFL counted libraries, we generally observe higher discovery rates at all percentages of the full counted libraries, and even more so when the 3’DGE counted libraries are initially subsampled to full SFL counted library sizes (Figure 5B), demonstrating that SFL shows improvements independent of the mapping rate. This result confirms previous reports showing that increasing read length above 50-bp does not improve read quantification (Chhangawala et al., 2015). Furthermore, similar results have been reported even when the same sequencing platform is used. A recent study reported a greater number of genes detected, as well as higher differential analysis discovery rates, in conventional RNA-seq than in 3’DGE at identical counted library sizes, using the Illumina HiSeq 2500 platform to generate both libraries (Xiong et al., 2017).

In summary, in this study we observe higher performance of SFL than 3’DGE, as measured by coverage, signal-to-noise, and biological recapitulation of known signal, with the performance of SFL often matching that of well-established “gold standards” (full coverage RNA-seq or microarrays). On the other hand, the fact that 3’DGE is shown to allocate a large number of reads to relatively fewer, highly expressed genes, makes this platform more suitable for problems where high accuracy in the differential quantification of highly expressed genes is needed. Furthermore, the ready availability of 3’DGE as a core-provided option, which allows for the out-sourcing of library preparation, sequence read pre-processing and gene quantification, is an additional value-added of the platform. Ultimately, the best-suited platform for a specific project will depend on the study goals, design, and availability of different resources. We believe our study presents useful results to make a more informed choice.

The utility of highly multiplexed RNA-seq crucially depends on the trade-off between cost and data quality, and on the nature of the experiments for which the platform would be ideally suitable. These will in general be experiments where the marginal information content of a single profile is relatively low, and thus justifies trading-off some data quality for reduced cost.

## Acknowledgment

Dr. Alexander A. Shishkin for his feedback during the development of the SFL protocol. SCRB-Seq libraries were prepared by the Broad Technology Labs and sequenced by the Broad Genomics Platform.

## Supplementary Material

Supplementary data and figures are available in the file, SupplementaryMaterial.pdf.

