## Supplementary Material for "Assessment of a highly multiplexed RNA sequencing platform and comparison to existing high-throughput gene expression profiling techniques"

### Supplementary Figures

**Table S1: Minimum within treatment group correlation of normalized gene expression across all genes**

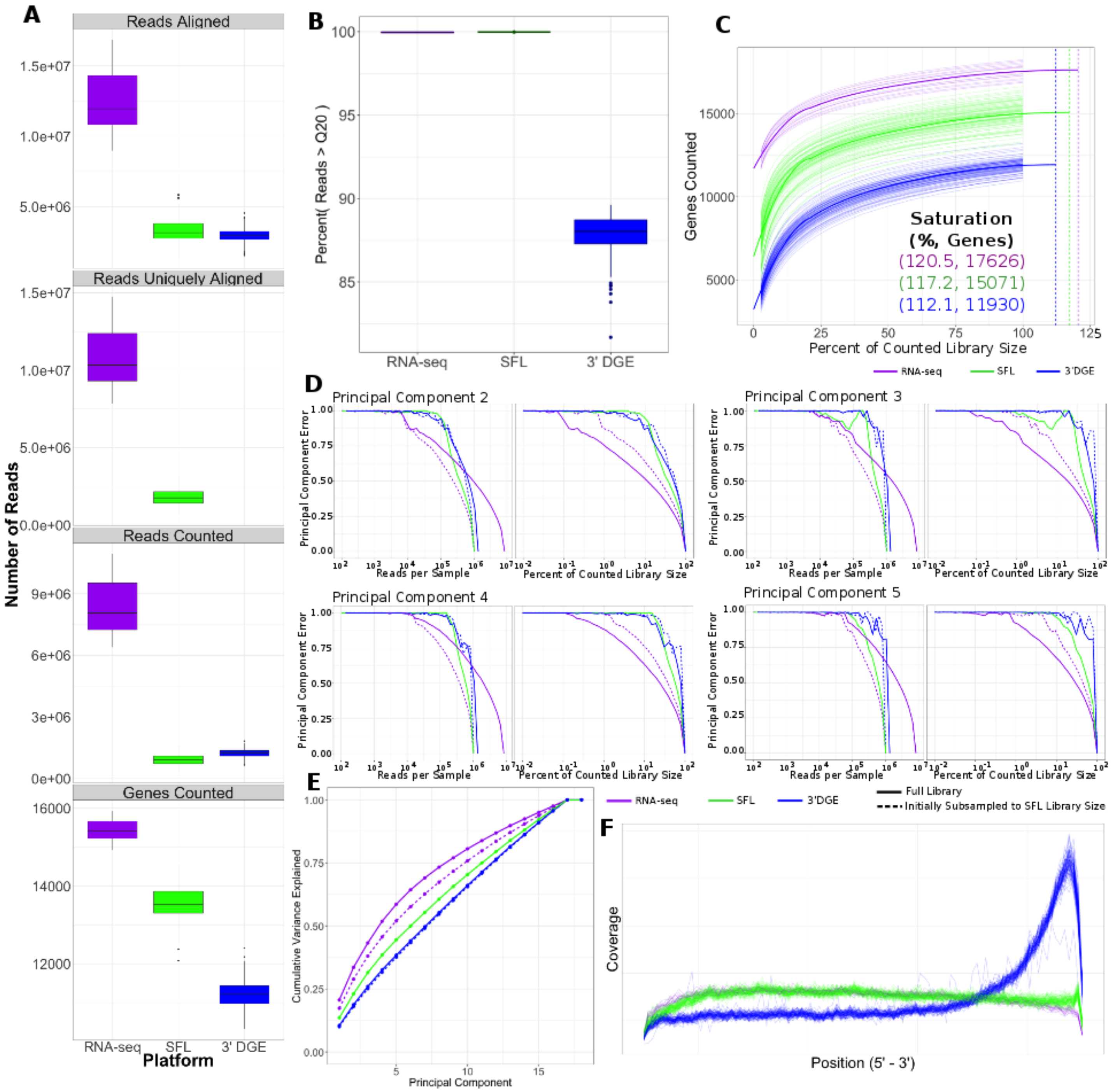

**Figure S1: Additional coverage analyses comparing Full Coverage RNA-seq, SFL, and 3'DGE**

A) Distribution of read counts at different preprocessing steps, as well as the number of counted genes.

B) Per platform boxplots of the percentage of reads in each sample with Phred Values > 20 (Q20).

C) Saturation analysis showing the number of genes with counts > 0 versus the percent subsampling of the full counted library size. Thick lines show the loess fit of each platform. Vertical lines show the estimated point of saturation, i.e. the minimum percentage of the full counted library at which the maximum number of genes are counted. This value is also given, as well as the estimated maximum number of counted genes.

D) Principal component error for PC's 2-5 for each platform, including subsampled full coverage RNA-seq and 3' DGE to the SFL library size.
E) Cumulative variance explained by each successive principal component.
F) Relative coverage of reads along transcripts from 5' to 3'.

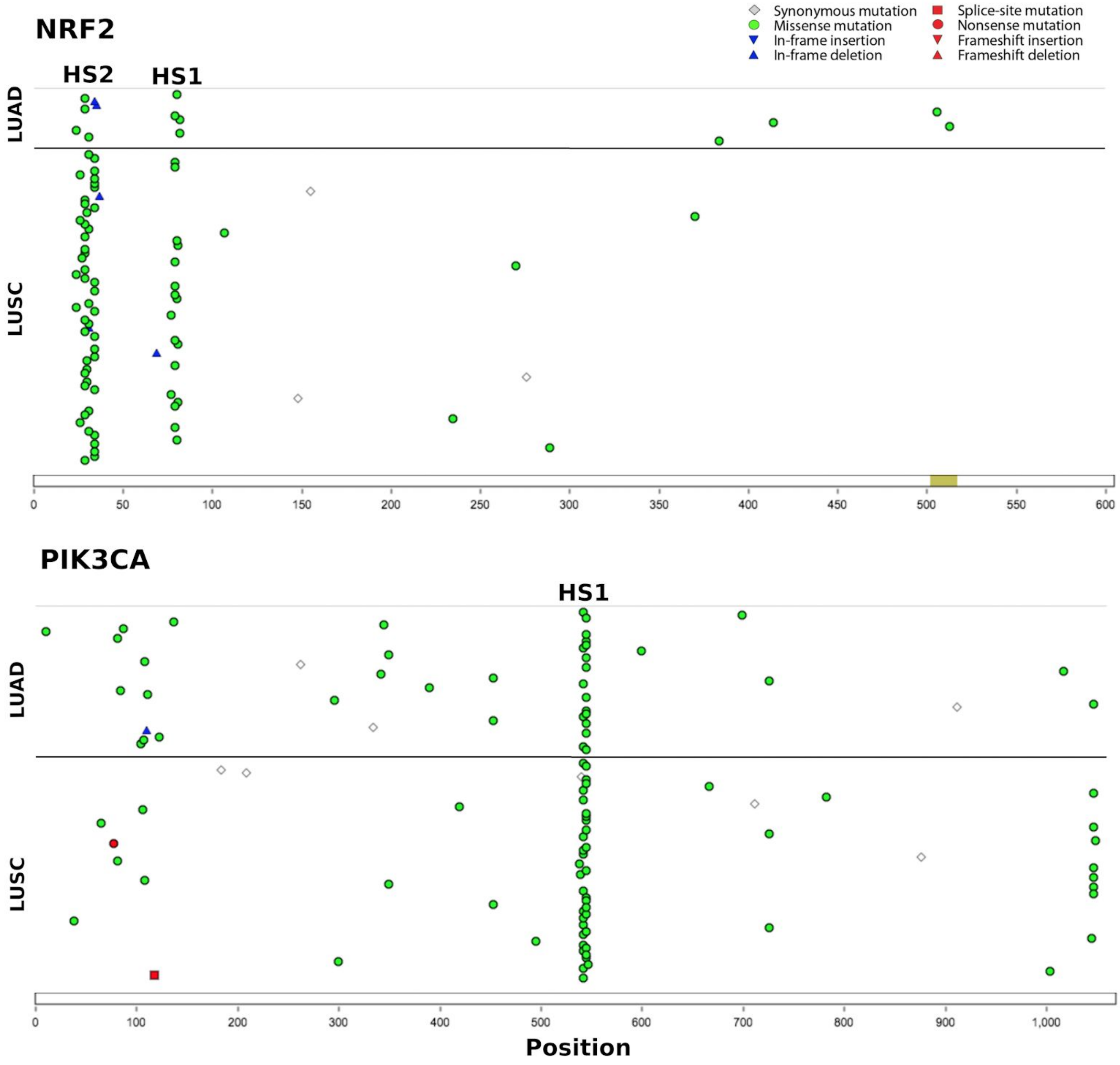

**Figure S2: Locations of mutations hotspots in NRF2 and PIK3CA**

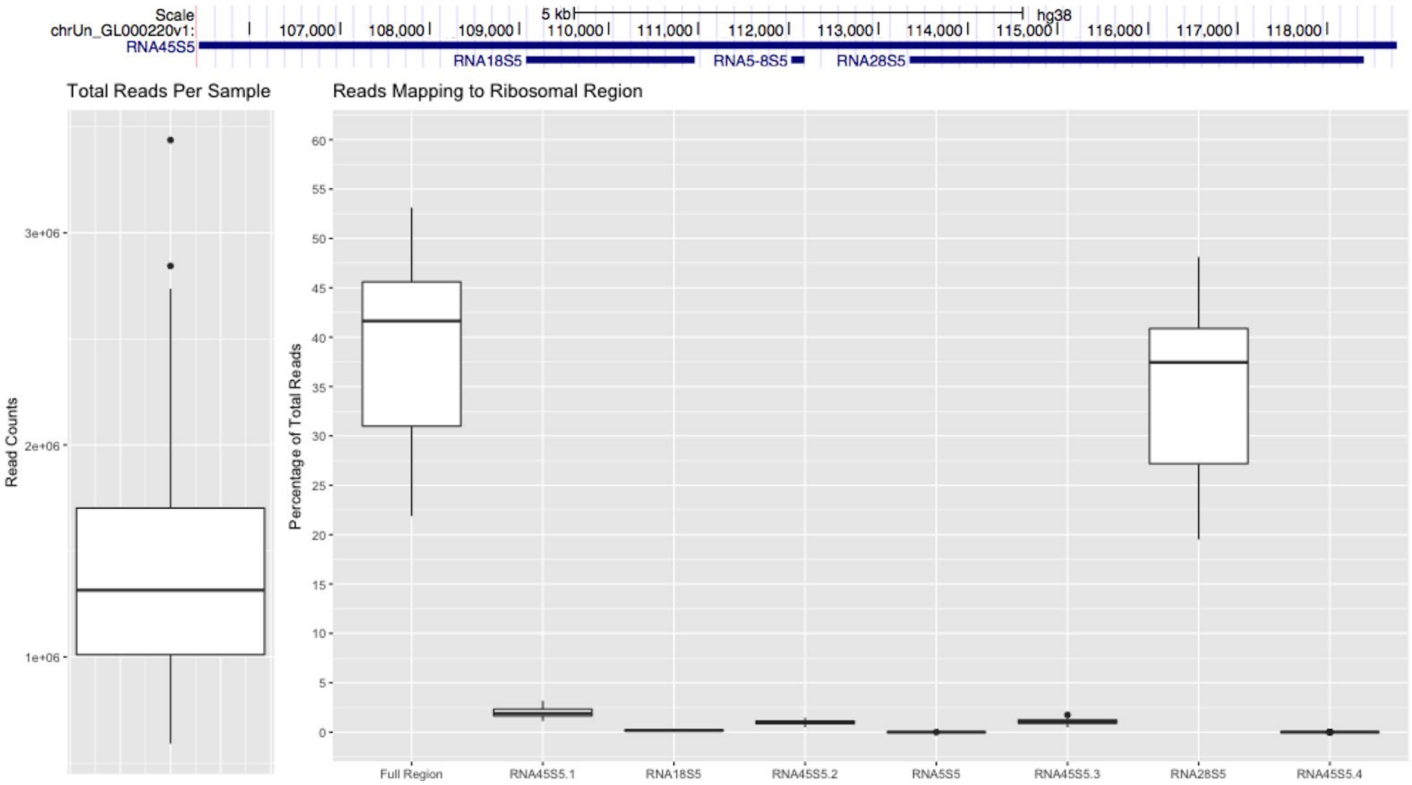

**Figure S3: Summary of rRNA contamination in SFL libraries**

The total library sizes for SFL samples (left) and proportion of these reads that align to a ribosomal region (RNA45S5) in the genome. The majority of these reads align to RNA28S5.

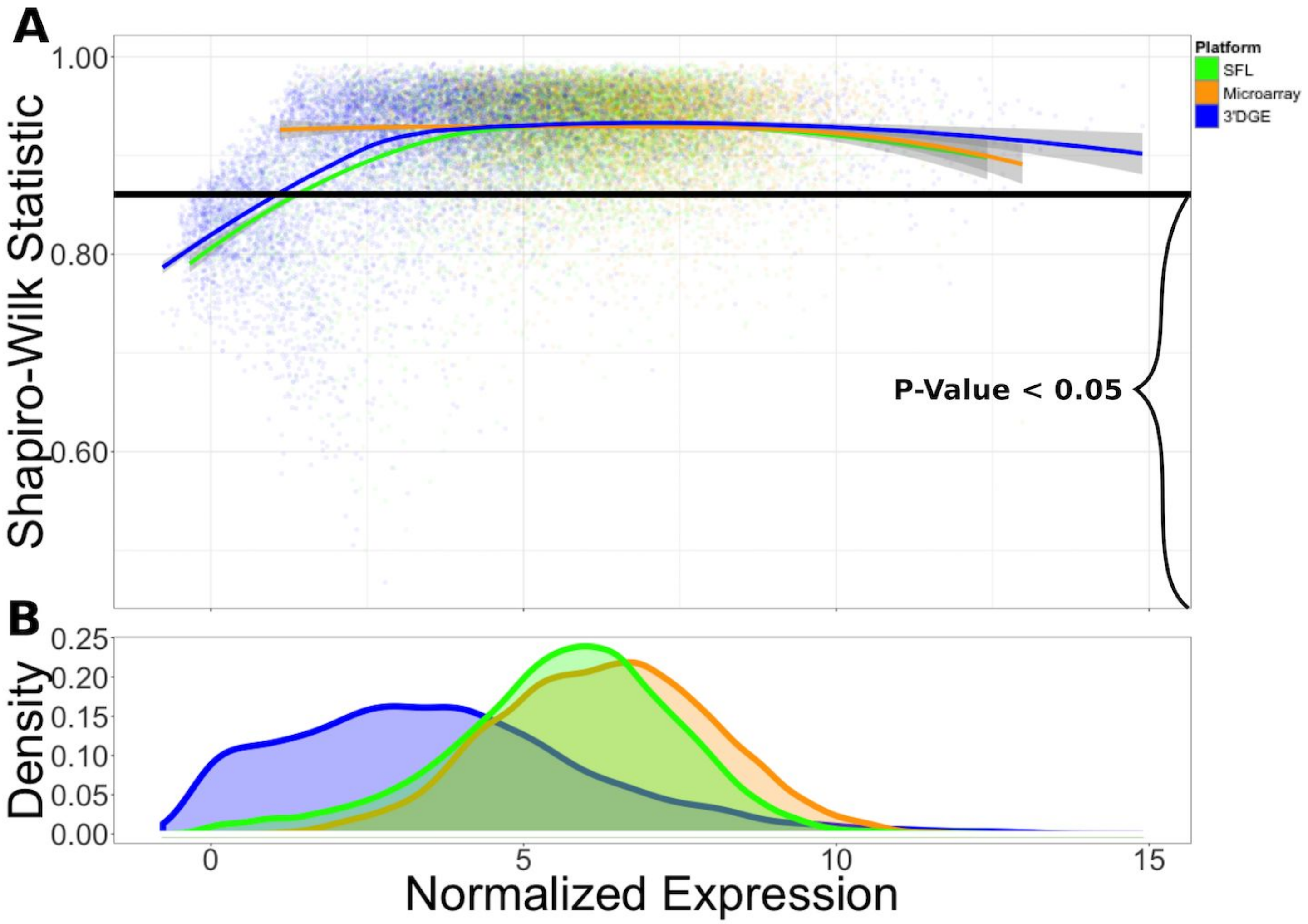

**Figure S4: Shapiro-Wilk Test Statistic VS Normalized Expression for SFL, Microarray, and 3' DGE**

A) Loess fit of the Shapiro-Wilks test statistic vs normalized expression across each platform. Values beneath the vertical black line are indicative of test statistics associated with nominal p-values < 0.05.

B) Distribution of mean normalized expression across all three platforms.

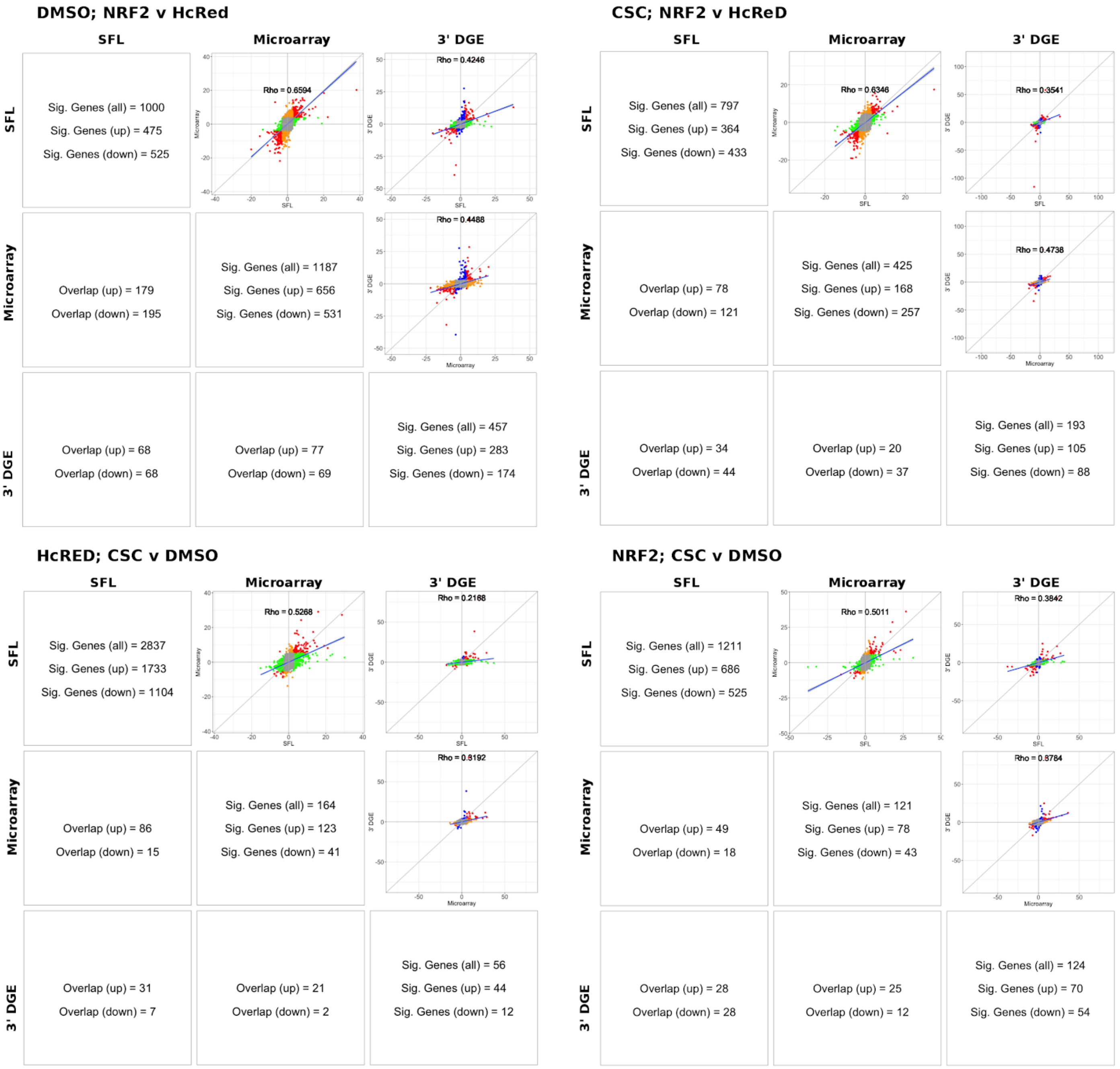

**Figure S5: Additional Differential Analysis Results (SFL; Microarray; 3'DGE)**

Comparison of gene discovery (FDR Q-Value < 0.05) by differential analysis with limma, comparing normalized gene expression, including the raw discovery rates, discovered gene overlap, and linear fits, comparing test statistics from each platform. Euler diagrams of these results are shown in Figure S6.

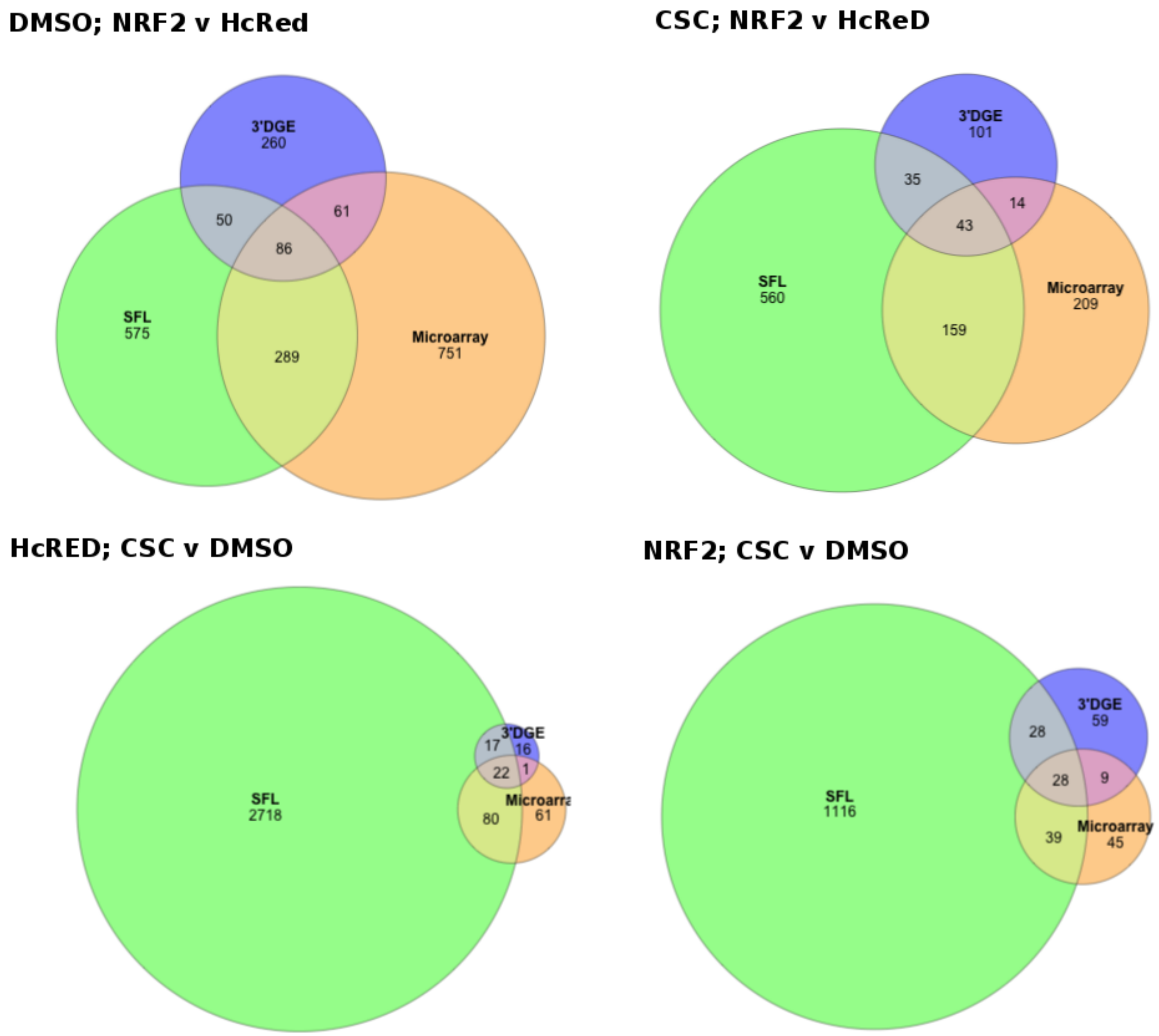

**Figure S6: Venn diagrams of gene discovery from differential analysis (SFL; Microarray; 3'DGE)**
Comparison of gene discovery (FDR Q-Value < 0.05) by two-group differential analysis with LIMMA.

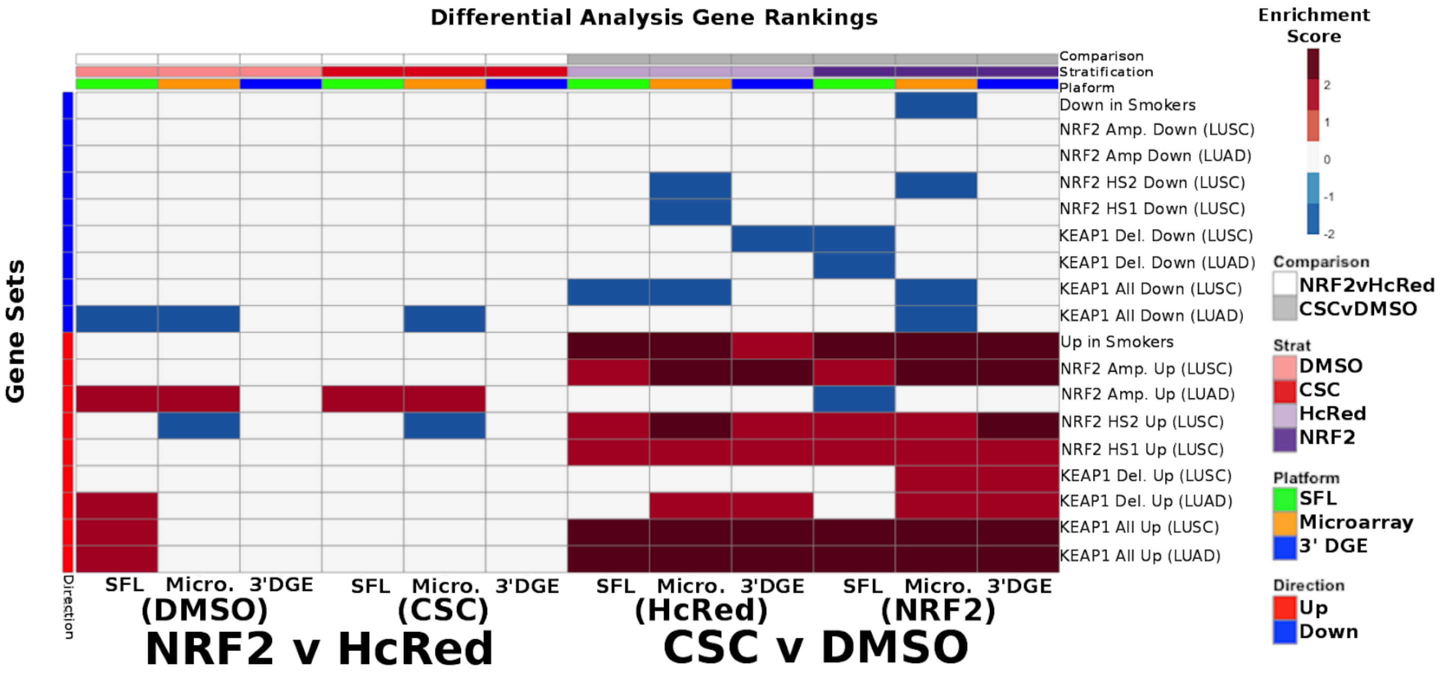

**Figure S7: Gene Set specific results of Smoking and Gene Mutation Signatures across SFL, 3'DGE and Microarray**

Gene-set enrichment results with respect to genotypic perturbations (left) and chemical perturbations (right) differential signatures across like samples within SFL, Microarray, and 3' DGE. Columns correspond to differential signatures comparing genotypic or chemical perturbation groups, stratified by a single chemical or genotypic perturbation group, respectively, e.g. the left-most column shows the enrichment results with respect to the "DMSO-treated; NRF2 *vs.* HcRed" signature within the samples (*stratum*) in SFL data. Row labels for the TCGA-derived genes sets follow the nomenclature "<GENE> <Mutation Type> <Direction>(<Source>)". Mutation type can be one of: *Amp.* for copy number gain, *Del.* for copy number loss, *HS#* for point mutations in hotspots (See Figure S2), and *All* for collapsed point mutations across the full length of the genes.

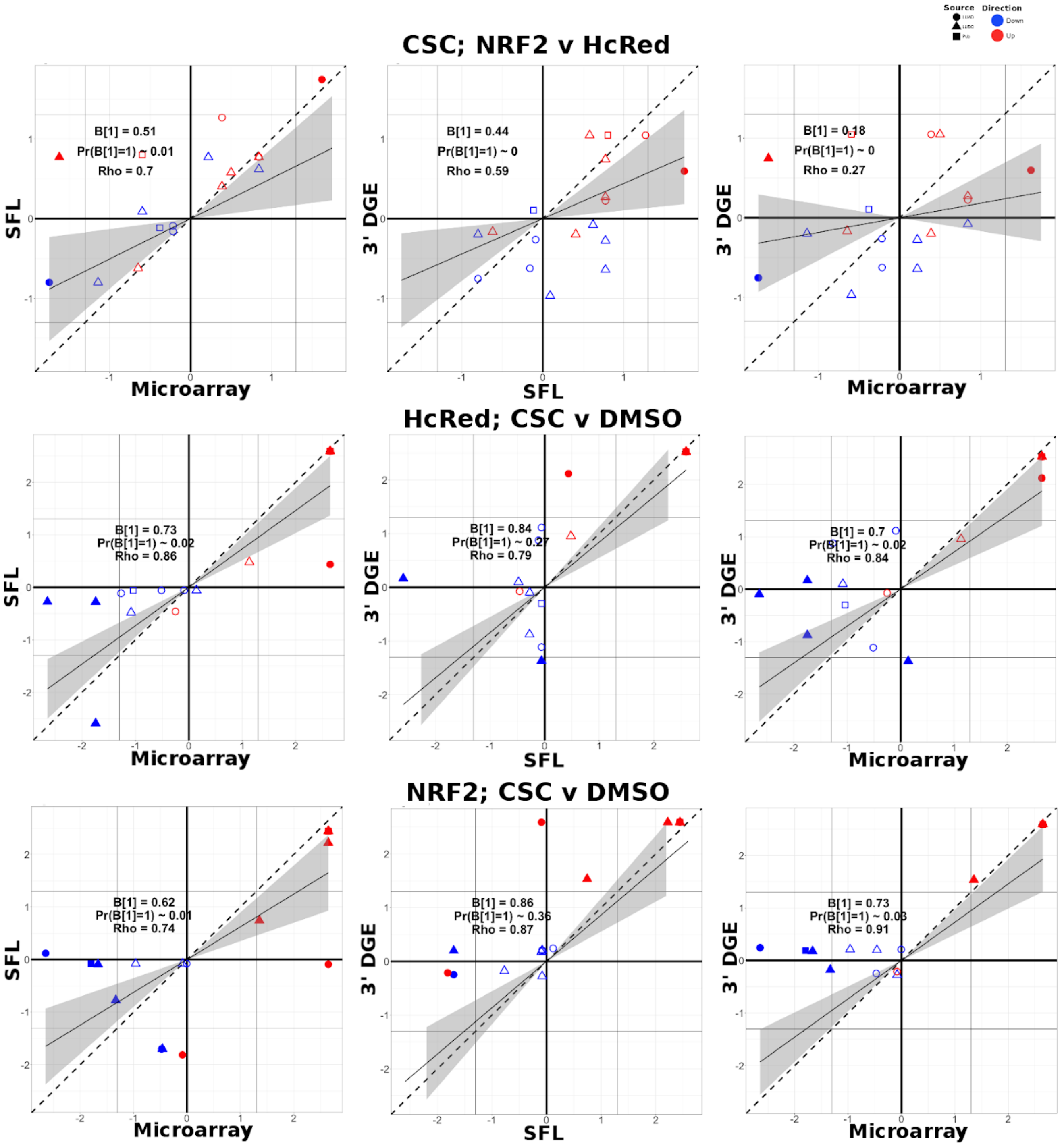

**Figure S8: Additional Biological Recapitulation Comparisons (SFL; Microarray; 3'DGE)**
Comparison of the gene set enrichment results between SFL, microarray and 3' DGE with respect to the "CSC-treated; NRF2 *vs.* HcRed", "HcRed-treated; CSC *vs.* DMSO", and "NRF2-treated; CSC *vs.* DMSO" differential signature. Shown are the transformed FDR q-values of the TCGA-derived gene sets corresponding to mutations of comparable gene sets. The |-Log10(FDR Q-values)| = 0.05 significance thresholds are shown as vertical and horizontal gray lines for the y and x-axes, respectively. Points of gene sets whose enrichment meets this threshold in either of the two platforms are filled in. Colors and shape of points denote direction and source of the gene set, respectively.

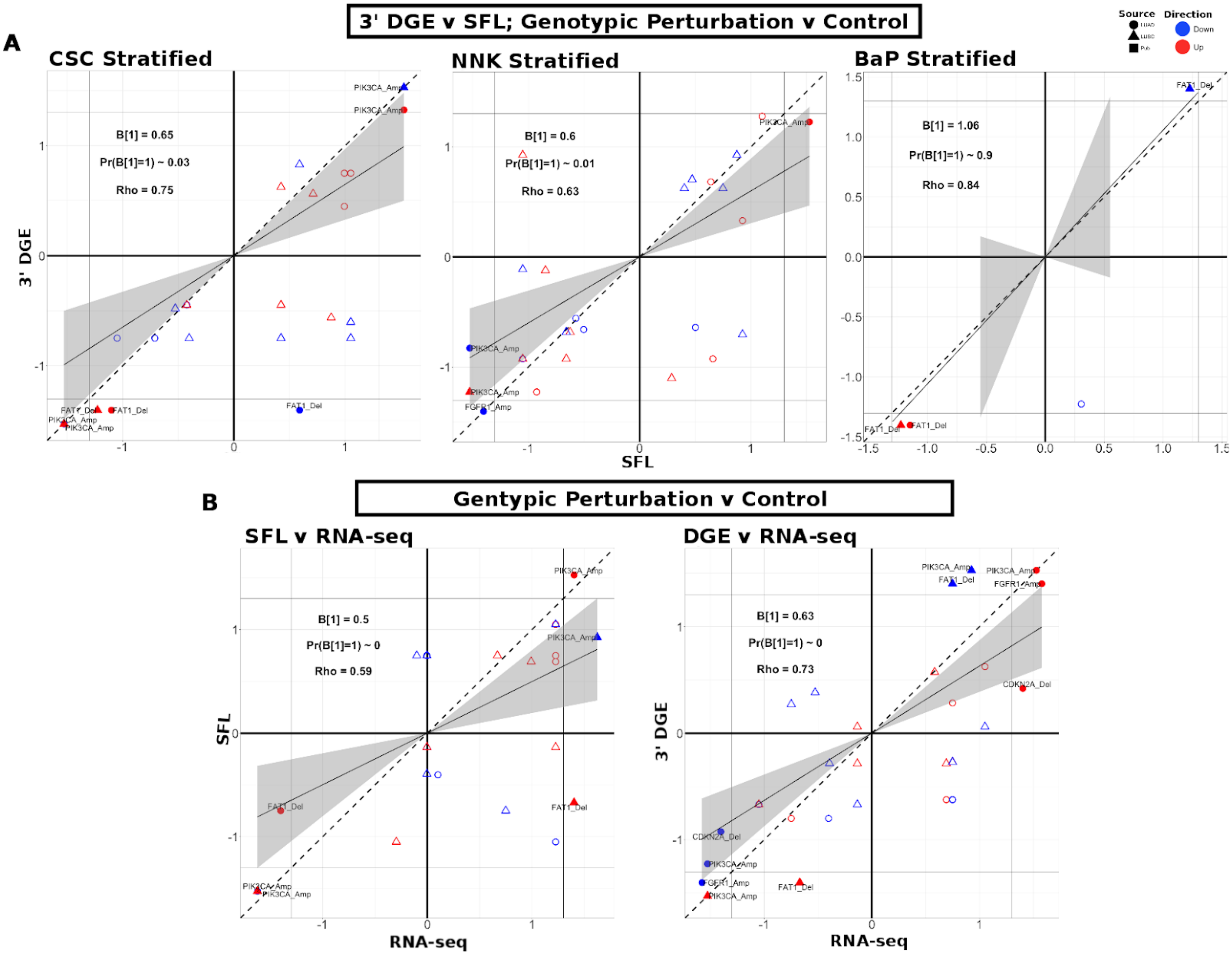

**Figure S9: Additional Biological Recapitulation Comparisons (RNA-seq; SFL; 3'DGE)**

A) Comparison of the gene set enrichment results between SFL and 3' DGE results for genotypic perturbation v control, stratified for BaP chemical exposure. Points indicate gene set comparisons to concordant signatures, derived from TCGA. In this case only FAT1 gene sets are shown because only FAT1 and GFP genotypic perturbations are available for BaP exposed samples. Shown are the transformed FDR Q-values from permutations based testing from preranked GSEA. The |-Log10(FDR Q-values)| = 0.05 significance thresholds are shown as vertical and horizontal gray lines for the y and x-axes, respectively. The names of these gene sets that meet this threshold in either of the two signatures are shown and their points are filled in. Colors and shape of points are indicative of the direction and source of the gene set, respectively.

B) Comparison of the gene set enrichment results between full coverage RNA-seq and either SFL or 3' DGE results for genotypic perturbation v control, stratified for DMSO exposures. Points indicate gene set comparisons to concordant signatures, derived from TCGA, e.g. PIK3CA mutation and CNA gene sets to PIK3CA vs HcRED preranked gene signatures. Shown are the transformed FDR q-values from permutations based testing from preranked GSEA. The |-Log10(FDR Q-values)| = 0.05 significance thresholds are shown as vertical and horizontal gray lines for the y and x-axes, respectively. The names of these gene sets that meet this threshold in either of the two signatures are shown and their points are filled in. Colors and shape of points are indicative of the direction and source of the gene set, respectively.

### Supplementary Data

#### RNAtag-seq Protocol (SFL) for batch of 96 RNA samples

##### Reagents:

- TURBO™ DNase, Life Technologies/Thermo Fisher, Cat.# AM2238
- FastAP Thermosensitive Alkaline Phosphatase (1 U/µL), Life Technologies/Thermo Fisher, Cat.# EF0651
- Buffer RLT (220 ml), Qiagen, Cat.# 79216
- T4 RNA Ligase 1 (ssRNA Ligase), High Conc. (30U/µL), NEB, Cat.# M0437M, and comes with the following reagents:
  - ATP (100mM) (aliquot and store at - 80$^o$C, always use fresh aliquot)
  - PEG 8000, 50% in water, 10X ligase buffer
- DMSO (Dimethyl sulfoxide, Hybri-Max), sterile, BioReagent, ⩾99.7%; Sigma, Cat.# D2650-5X10ML
- RNase Inhibitor, Murine (40U/µL); NEB, Cat.# M0314L (15,000 units)
- T4 Polynucleotide Kinase, NEB, Cat.# M0201L
- NEBNext® Q5® Hot Start HiFi PCR Master Mix, NEB, Cat.# M0543S
- AffinityScript Multiple Temperature cDNA Synthesis Kit, 50 reaction; Agilent, Cat.# 200436 – includes the dNTPs, 10x RT buffer, RNase Block Ribonuclease Inhibitor (40U/µL)
- Dynabeads® MyOne™ Silane, Life Technologies/Thermo Fisher, Cat.# 37002D
- ExoSAP-IT PCR Product Cleanup, Affymetrix, Cat.# 78200
- ZR-96 RNA Clean & Concentrator, Zymo Research, Cat.# R1080
- RNA Clean & Concentrator™-5 columns, Zymo Research, Cat.# R1015
- RNAClean XP beads, Agencourt/Beckman, Cat.# A63987 (40mL)
- Ribo-Zero™ Magnetic Gold Kit (Human/Mouse/Rat); Epicentre/Illumina, Cat.# MRZG126
- Sodium hydroxide solution, volumetric, 4 M NaOH (4N), Sigma, Cat.# 35274-1L
- Hydrochloric acid, 36.5-38.0%, BioReagent, for molecular biology, Sigma, Cat.# H1758-100ML
- Nuclease-Free Water (not DEPC-Treated), Life Technologies/Thermo Fisher, Cat.# AM9930
- TE Buffer, 1X Solution pH 8.0, Low EDTA, Affymetrix, Cat.# 75793 500 ML
- Agilent RNA 6000 Pico Kit, Agilent, Cat.# 5067-1513
- High Sensitivity DNA Kit, Agilent, Cat.# 5067-4626
- Invitrogen™ UltraPure™ 0.5M EDTA, pH 8.0, Fisher Scientific/Thermo Fisher, Cat.# 15-575-020

##### Procedure:

1. **Quality and quantity of RNA with Agilent BioAnalyzer**
   1) Check RNA quality by running on the Agilent BioAnalyzer
   2) Place 200 ng of total RNA in one well of the 96-well PCR plate
   3) Increase the volume to 18 µL with nuclease-free water
2. **FastAP + PNK treatment of RNA for large fragments (when library will be sequenced 76-index-76 or 100-index-100):**

   **2.1. Fragmentation using 2x FastAP buffer**

   Set up the following reaction:

| Reagents | 1 reaction | Final Conc. |
|---|---|---|
| FastAP buffer (10x) | 2 µL | 2x |
| RNA (from step 1) | 18 µL | 200ng total RNA |
| Total volume (µL) | 20 µL | |

1) Add 2 µL of FastAP buffer to the RNA and mix well.
2) Incubate on thermal cycler for 3 minutes at 94°C
   Note: Fragmentation time can vary from 90 seconds to 4 minutes, depending on RNA sizes you will need (or if RNA is partially degraded)
3) Quickly place on ice

**2.2. FastAP treatment**

Set up the following reaction mix (on ice)

| **Repair mix** | **1 well/tube** | | **x 96 samples** (10% overhang) |
|---|---|---|---|
| Fragmented RNA (in tube/plate on ice) | ***20 µl*** | | |
| ***Add master-mix*** | | | |
| $H_2O$ | 21 µl | *30 µl* | 2217.6 µl |
| TURBO™ DNase | 2 µl | | 211.2 µl |
| 10× FastAP Buffer | 3 µl | | 316.80 µl |
| RNase Inhibitor, Murine ( NEB, M0314L) | 0.5 µl | | 52.80 µl |
| FastAP enzyme | 3.5 µl | | 369.60 µl |
| ***Total*** | ***50 µl*** | | |

1) Directly add 30 µL of master-mix into each well/tube with the fragmented RNA (on ice), mix well
2) Incubate for 12 minutes at 37°C on a thermoshaker

**2.3. PNK treatment**

**(**After 12 min, add 150 µl of T4 PNK master-mix directly into FastAP-treated RNA):

| **Fragmentation/Repair (mix)** (room temperature) | **1 well/tube** | | **x 96 samples** (10% overhang) |
|---|---|---|---|
| FastAP-treated RNA | ***50 µl*** | | |
| ***Add* T4 PNK *master-mix into FastAP mix*** | | | |
| $H_2O$ | 114 µl | *150 µl* | 12038.4 µl |
| TURBO™ DNase | 2 µl | | 211.2 µl |
| 10× T4 PNK buffer | 20 µl | | 2,112 µl |
| RNase inhibitor | 3.5 µl | | 369.60 µl |
| T4 Polynucleotide Kinase (NEB, M0201L) | 10.5 µl | | 1,108.80 µl |
| ***Total*** | ***200 µl*** | | |

1) Mix well after adding master-mix
2) Incubate for 30 minutes at 37°C on a thermoshaker

**2.4. ZR-96 RNA Clean & Concentrator to clean up PNK-treated RNA**

1) Prepare adjusted RNA Binding Buffer (as needed). Mix an equal volume of buffer and ethanol (95-100%). [Example: Mix 200 µl buffer and 200 µl ethanol.]
2) Add 2 volumes of the adjusted buffer to the sample and mix. [Example: Mix 400 µl adjusted buffer and 200 µl sample.]
3) Transfer the sample to each well of a Zymo-Spin™ I-96 Plate1 mounted on a Collection Plate and centrifuge for 5 minutes. Re-load the flow-through to the same well of Zymo-Spin™ I-96 Plate1 mounted on a Collection Plate and centrifuge for 5 minutes again, and then discard the flow-through.
4) Add 400 µl/well RNA Prep Buffer and centrifuge for 5 minutes. Discard the flow-through.
5) Add 700 µl/well RNA Wash Buffer and centrifuge for 5 minutes. Discard the flow-through.

6) Add 400 µl/well RNA Wash Buffer and centrifuge for 5 minutes to ensure complete removal of the wash buffer. Mount the plate carefully onto an Elution Plate.
7) Add ≥10 µl/well DNase/RNase-Free Water directly to the matrix and centrifuge for 5 minutes. [To increase RNA yield, *put first flow-through again on the same column, and re-elute with the same 12µl of water.*] (Final volume may be 8-10µl)
   The eluted RNA can be used immediately for library prep (first ligation) or stored at -80 C. Use the Cover Foil to prevent evaporation.
8) Optional: use 1µl for RNA Pico chip and use about 7µL for library prep.

[All centrifugation steps should be performed at 3,000-5,000 x g.]

#### 3. FIRST LIGATION (RNA/RNA) 3' Linker Ligation

NOTE: *Set up RNA/barcoded adapter in single tube or use the TempPlate No-skirt PCR Plates for batched samples (these rigid plates are easy to handle and can mix well by flicking by hand) – with a razor, cut a column (for 8 samples) or row (for 12 samples) and use as strip and cover with the Flat PCR 12-cap strips (these strips fit tightly on these plates and will not leak)*

1) Add 1 µL of barcoded RNA oligo/adapter (40 pmoles, see Appendix for sequence information) to 6.5 µL of FastAP & PNK-treated (dephosphorylated/repaired) RNA:

| | | |
|---|---|---|
| **Repaired RNA , 50 ng** | 6.5 µl | 65 C, 2 min → ice |
| Barcoded RNA adapter (40 µM), 40 pmoles *[Note: for 100ng+ of starting RNA use 40-100pmoles of adapters, for <100ng use 20 pmoles]* | 1 µl | |
| 100% DMSO | 1.5 µl | |
| Total RNA part | 8.5 µl | *(0.5 µl will evaporate)* |

2) Denature RNA when master mix is almost ready: heat RNA samples (RNA + adapter + DMSO) at 65$^{o}$C for 2 minutes → place on ice
3) Set up ligation mix below
   **NOTE:**
   - Make up ligation mix at room temp so the reagents don't start precipitating when combined (*if DMSO is added directly into cold buffer it will precipitate*)
   - Pipette very slowly for accurate aspiration of PEG (very viscous)
   - When setting up mix for multiple reactions include 25% extra to account for pipetting
   - All reagents except enzymes (-20 $^{o}$C ) should be stored at -80 $^{o}$C in single use aliquots:

| **Ligation Mix (set up at room temp)** | **1 Reaction** | **120 Reaction Mix (for a set of 96)** | **x 120** |
|---|---|---|---|
| H2O | 1.3 µL | 156 µL | 1380 µL |
| 10× NEB ligase buffer | 2 µL | 240 µL | |
| DMSO (100%) | 0.5 µL | 60 µL | |
| ATP (100mM - stored at -80) **Use fresh ATP!!!** | 0.2 µL | 24 µL | |
| PEG8000 (50% stock) | 6 µL | 720 µL | |
| RNase inhibitor, Murine (40U/µL) (NEB) | 0.3 µL | 36 µL | |
| T4 RNA ligase 1, Hi Conc. (30U/µl), 36 Units (NEB) | 1.2 µL | 144 µL | |
| **Total (RNA + ligation mix) REACTION** | **20 µL** | | |

4) Mix really well by flicking tube since the solution is very viscous
5) Quick spin of master-mix tube
6) Add 11.5 µL of ligation mix to each well containing denatured RNA (~8.5 µL)

7) Mix well MANY times; mix by flicking since the solution is very viscous
(If setting this up on a robot, the reaction should be mixed up/down for 5-10 minutes with low retention tips to mix well!)
8) Incubate at 23°C (room temp) for 1 hour 15 minutes [shake well IN HANDS every 15-30 minutes!!!]

**4. Pooling step: Zymo RNA Clean & Concentrator™-5 column**

[Maximum binding capacity of columns is 5 µg; do not exceed when pooling samples]

1) Prepare adjusted RNA Binding Buffer (as needed). Mix an equal volume of buffer and ethanol (95-100%). [Example: Mix 200 µl buffer and 200 µl ethanol.]
2) Add 2 volumes of the adjusted buffer to the ligation mix from above and mix well (use low retention tips and re-suspend well) [Example: Mix 40 µl adjusted buffer and 20 µl ligation mix.], pool 12 samples together (or other numbers as planned)
3) Transfer the pooled sample to the Zymo-Spin™ IC Column in a Collection Tube (may have to load several times if pooled sample volume > 700 µl) and centrifuge for 30 seconds. Re-load the flow-through to the same Zymo-Spin™ IC Column in a Collection Tube and centrifuge for 30 seconds again, and then discard the flow-through.
4) Add 400 µl RNA Prep Buffer to the column and centrifuge for 30 seconds. Discard the flow-through.
5) Add 700 µl RNA Wash Buffer to the column and centrifuge for 30 seconds. Discard the flow-through.
6) Add 400 µl RNA Wash Buffer to the column and centrifuge for 2 minutes to ensure complete removal of the wash buffer. Transfer the column carefully into an RNase-free tube.
7) Add 16 µl DNase/RNase-Free Water directly to the column matrix and centrifuge for 30 seconds, re-elute with the flow-through. The eluted RNA can be used immediately or stored at -80 $^{o}$C.

Optional: Save 2 µL for QC - Run on Agilent RNA Pico chip to check the fragmentation profile

[All centrifugation steps should be performed at 10,000 - 16,000 x g.]

**5. Quick Protocol for Ribo-Zero™ rRNA Depletion**

(For details, please see Illumina "Ribo-Zero Magnetic Kits User Guide 15066012A")

**5.1 <u>Wash the Magnetic Beads</u>**

The Magnetic Beads must be washed before using Ribo-Zero Kits. It is important to use the beads at room temperature.

**IMPORTANT: Do not freeze or place the Magnetic Beads on ice. Damaged (frozen) beads will impact kit performance.**

**Procedure:**

Each Ribo-Zero reaction requires 225 µl of the Magnetic Beads. Up to 1,350 µl of the Magnetic Beads (sufficient for 6 Ribo-Zero reactions) can be washed in a 1.5 ml tube. Use a 15 ml tube if batch-washing more than 1,350 µl of beads.

1) Remove the Magnetic Core Kit components from storage at 2°C to 8°C and allow them to equilibrate to room temperature. Do not place the Magnetic Core Kit or Magnetic Beads on ice.

2) Vigorously vortex the Magnetic Beads to homogeneity.

3) Pipette the Magnetic Bead suspension slowly to avoid air bubbles and to ensure pipetting the correct volume.

4) Place the tube containing the Magnetic Beads on the magnetic stand for at least 1 minute until the solution appears clear.

5) With the tube still on the stand, remove and discard the supernatant.
[CAUTION: The supernatant contains 0.1% sodium azide.]

6) Remove the tube from the stand and add an equal volume of RNase-Free Water. Vigorously vortex the Magnetic Beads.

7) Repeat steps 4, 5, and 6 (i.e., wash the beads a total of 2 times with RNase-Free Water).
8) After the second water wash, place the tube in a magnetic stand for 1 minute, then remove and discard the water.
9) Remove the tube from the magnetic stand. Add a volume of Magnetic Bead Resuspension Solution equal to the number of reactions × 60 µl (e.g., for 6 reactions, add 6 × 60 µl = 360 µl Magnetic Bead Resuspension Solution). Mix well by vigorous vortexing.
[NOTE: The volumes of the beads and Resuspension Solution are additive. Although the washed beads are resuspended in 60 µl per reaction, each reaction uses 65 µl of resuspended beads.]
10) Aliquot 65 µl of the washed and resuspended Magnetic Beads into new 1.5 ml RNase-free microcentrifuge tubes corresponding to the number of Ribo-Zero reactions.
11) Optional: Add 1 µl of RiboGuard RNase Inhibitor to each tube of resuspended Magnetic Beads, and mix briefly by vortexing.
12) Store the washed Magnetic Beads at room temperature until required in "Remove rRNA" step. Return the remaining Magnetic Core Kit components to storage at 2°C to 8°C.

#### 5.2 Treat the Sample with Ribo-Zero rRNA Removal Solution

In this step, the Ribo-Zero rRNA Removal Solution (probes) hybridizes to the rRNA in the sample. RNA samples must be DNase-treated and purified prior to treatment with Ribo-Zero rRNA Removal Solution.

CAUTION: The maximum volume of the RNA sample and the volume of the Ribo-Zero rRNA Removal Solution used per reaction depends on the amount of total RNA in the sample (see Table below).

| Amount of Input Total RNA for All Ribo-Zero Kits (Except Epidemiology) | Maximum Volume of Total RNA That Can Be Added to Each Reaction | Volume of Ribo-Zero rRNA Removal Solution Used per Reaction |
|---|---|---|
| 1–2.5 µg | 28 µl | 8 µl |
| > 2.5–5 µg | 26 µl | 10 µl |

**Procedure:**

1) Remove Ribo-Zero rRNA Kit components from storage at -65°C to -80°C and allow them to equilibrate to room temperature.
2) For each sample, add the following reagents in a 0.2 ml or 0.5 ml RNase-free microcentrifuge tube. Combine in the order given:

| **Reagent** | **Volume (µl)** |
|---|---|
| RNase-Free Water | x |
| Ribo-Zero rRNA Reaction Buffer | 4 |
| RNA sample (see Table above) | y |
| Ribo-Zero Removal Solution (see Table above) | 8-10 |
| **Total Volume per Sample** | 40 |

3) Fully mix by pipetting 10–15 times. Incubate at 68°C for 10 minutes.
   - Return the remaining Ribo-Zero rRNA Removal Solution and Ribo-Zero rRNA Reaction Buffer to storage at -65°C to -80°C.
4) Remove the tubes from heat and centrifuge briefly to collect any condensation.
5) Incubate the tubes at room temperature for 5 minutes.

#### 5.3 Remove rRNA

This step is the most important part of the Ribo-Zero procedure. It combines probe-hybridized samples (from the previous step "Treat the Sample with Ribo-Zero rRNA Removal Solution") with the washed,

room temperature Magnetic Beads (from the "Wash the Magnetic Beads" step).
[CAUTION: The washed Magnetic Beads must be at room temperature for use in this step. The order of this addition is critical and can impact rRNA removal efficiency if done incorrectly!!!]

1) Transfer the probe-hybridized RNA sample to the washed, room temperature Magnetic Beads.

| **Reagent** | **Volume (µl)** |
|---|---|
| Probe-hybridized RNA sample | 40 |
| Washed, room temperature Magnetic Beads | 65 |
| **Total Volume per Sample** | 105 |

Without changing the pipette tip, immediately and thoroughly mix the contents of the tube by pipetting 10–15 times, as shown in Figure below. Set the tube aside at room temperature and repeat step 1 for each sample.

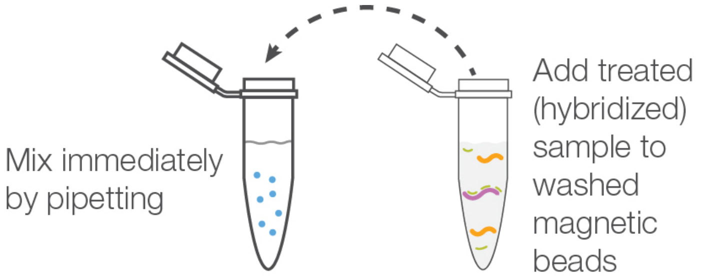

2) Cap the tubes and vortex at high speed for a minimum of 10 seconds. Avoid vortexing the beads/sample into the tube cap. Perform this step for each sample.
3) Incubate the tubes at room temperature for 5 minutes.
4) Incubate the tubes at 50°C for 5 minutes.
5) Remove the tubes from heat and immediately place them on a magnetic stand for at least 1 minute until the solution appears clear.
6) Carefully remove the supernatant (85–90 µl) containing the sample depleted of rRNA, and transfer to an appropriately sized RNase-free microcentrifuge tube.
[CAUTION: The supernatant contains the depleted RNA sample.]
[NOTE: Remove all the Magnetic Beads from the sample. The Magnetic Beads have the unwanted rRNA and Ribo-Zero probes bound to them. If any Magnetic Beads are still visible in the supernatant, place the collected supernatant onto the magnetic stand for 1 minute. Remove the supernatant containing the depleted RNA sample and transfer it to a new RNase-free tube.]
7) Place the supernatant (sample depleted of rRNA) on ice and proceed to the next step "Purify the Ribo-Zero–Treated RNA". Alternatively, keep the supernatant at -20°C overnight or at -65°C to -80°C for long-term storage.

**<u>5.4 Purify the Ribo-Zero–Treated RNA using Zymo RNA Clean & Concentrator-5 kit</u>**

[Maximum binding capacity of columns is 5 µg; do not exceed when pooling samples]

1) Prepare adjusted RNA Binding Buffer (as needed). Mix an equal volume of buffer and ethanol (95-100%). [Example: Mix 200 µl buffer and 200 µl ethanol.]
2) Add 2 volumes of the adjusted buffer to the rRNA-depleted RNA sample from previous step and mix well (use low retention tips and re-suspend well) [Example: Mix 180 µl adjusted buffer and 90 µl depleted RNA.]
3) Transfer the buffer-RNA mix to the Zymo-Spin™ IC Column in a Collection Tube and centrifuge for 30 seconds. Discard the flow-through.
4) Add 400 µl RNA Prep Buffer to the column and centrifuge for 30 seconds. Discard the flow-through.
5) Add 700 µl RNA Wash Buffer to the column and centrifuge for 30 seconds. Discard the flow-through.
6) Add 400 µl RNA Wash Buffer to the column and centrifuge for 2 minutes to ensure complete removal of the wash buffer. Transfer the column carefully into an RNase-free tube (not provided).

7) Add 12.5 µl DNase/RNase-Free Water directly to the column matrix and centrifuge for 30 seconds. [*Better bind on column twice (put first flow-through again on the same column), and elute twice with the same water. This will help improve recovery/yield of RNA.*]

The final elution should be about 11 µl. The eluted RNA can be used immediately or stored at -80 C.

[All centrifugation steps should be performed at 10,000 - 16,000 x g.]

**6. First Strand cDNA synthesis**

1) Take 11 µL rRNA-depleted RNA (use all the material from above)
2) Add 1 µL of AR2 RT primer (20 µM, see Appendix for sequence information)
3) Mix well
4) Heat the mixture to 65°C for 2 minutes and immediately place on ice
5) Make RT mix below: For multiple samples, a master mix can be prepared ahead of time and added to the RNA/AR2 tube on ice.

**Add (in order on ice):**

| **RT Mix** | **1 reaction** | **X 9 for 8 reactions (12.5% overhang)** |
|---|---|---|
| H2O | 2 µL | 18 µL |
| 10× AffinityScript RT Buffer | 2 µL | 18 µL |
| 100mM DTT | 2 µL | 18 µL |
| 25mM dNTP Mix | 0.8 µL | 7.2 µL |
| RNase inhibitor, murine (40U/µL) | 0.3 µL | 2.7 µL |
| AffinityScript RT Enzyme | 0.9 µL | 8.1 µL |
| Total (+ 12 µL rRNA depleted-RNA + primer) | 8→20 µL | |

6) Add 8 µL of RT mix to the 12µL rRNA depleted RNA + AR2 RT primer on ice
7) Mix well (shake in hands) and spin for 5 seconds
8) Incubate at 55°C for 45 minutes, then 4°C for 1 minutes

**7. RT primer removal after RT using ExoSAP-IT**

1) Add 4-5 µl (1 µl per 5 pmoles of RT primer – 1 µl 20 µM RT primer is equal to 20 pmoles) of ExoSAP-IT directly into 20 µl of RT reaction
2) Incubate at 37°C for 15 minutes

**8. RNA degradation after RT**

NOTE: make fresh working stock solutions of NaOH and Hydrochloric acid

1) Add 1 µl of 0.5M EDTA
2) Add 1 µl of 3M NaOH - incubate at 70°C for 12 minutes.
3) After incubation, neutralize with 1 µl of 3M HCL, mix well
4) Total volume = 27 µL

**9. RT primer Silane(Dynabeads® MyOne) cleanup**

1) Use 10 µl of beads/sample, add some RLT to the beads to rinse, remove supernatant.
2) 10µl of beads/sample + 50µl of RLT, mix well and on the magnet stand to remove supernatant. Re-suspend magnetic beads with RLT to 81µl.
3) Bind with 3.0× of fresh RLT (with beads) and 0.6× (Reaction mix + RLT volume) EtOH, mix well:

| | |
|---|---|
| Reaction mix | 27 µl |
| RLT with beads | 81 µl |
| *mix well for 1 min* | |
| EtOH | 65 µl |

| mix well for 2-5 min |
|---|

Magnet separation, remove supernatant
4) Wash beads in 123 µl of fresh 70% EtOH twice
5) Magnet separation, remove supernatant
6) Let air-dry at RT for 3-10 minutes (until beads stop shining, but do NOT over-dry them!)
7) Keep Silane beads, add 5.6 µl (0.6 µl will evaporate) of Adapter + DMSO solution to the beads [This is for the next step - "second ligation"]:

| | | |
|---|---|---|
| $H_2O$ + low TE mix (1V+1V) | 4 µl | *70°C, 2 min → cold ice* |
| 3Tr3 universal adapter, 80 pmoles (100 µM | 0.8 µl | |
| 100% DMSO | 0.8 µl | |
| *Total* | *5.6 µl* | *(0.6 µl will evaporate)* |

#### 10. Second LIGATION (ssDNA/ssDNA) 3' Linker Ligation with beads

(Use ultra low-retention tips!!! See Appendix for 3Tr3 adapter sequence information)

1) cDNA (contain beads) + 3Tr3 adapter + DMSO mix -> heat at 70°C for 2 minutes -> place on ice
2) Make ligation reaction mix: [For multiple samples, a master-mix can be prepared ahead of time and added to the cDNA/3Tr3 tube on ice. See notes for the first ligation about how to make master-mix.]

| 2nd Ligation Mix | 1 reaction |
|---|---|
| | 2.4 µl |
| 10× NEB Buffer | 2 µL |
| DMSO (100%) | 0.2 µL |
| ATP (100 mM) | 0.2 µL |
| PEG8000 (50% stock) | 9 µL |
| T4 RNA Ligase 1 (30 U/µL), 36 U | 1.2 µL |
| Total (+ 5 µl cDNA+adapter+DMSO mix) | 20 µL |

3) Swirl the cDNA/beads/water with pipet tip PRIOR to dispensing 15 µL ligation mix
4) Mix well by pipetting up and down ~10 times (pipetting MANY times, using ultra low-retention tips!!) or cap tubes and flick several times; solution is viscous
5) Quick spin (low speed centrifuge, to get everything to bottom of tube)
6) Incubate at 23°C for 2-4 hours or overnight

#### 11. Silane linker cleanup to remove the remaining adapters

1) Take extra 5 µl of Silane beads/sample, add some RLT to the beads to rinse, remove supernatant.
2) 5µl of beads/sample + 30µl of RLT, mix well and put on the magnet stand to remove supernatant, then re-suspend magnetic beads with RLT to 61µl
3) Bind with 3.0× of fresh RLT and 0.5× (Ligation mix + RLT volume) EtOH, mix well (e.g., pipetting up and down 15 times):

| | |
|---|---|
| Ligation mix (From above) | 20 µl |
| RLT with beads | 61 µl |

| | |
|---|---|
| *mix well for 1 min* | |
| EtOH | 40 µl |
| *mix well for 2 min* | |

Magnet separation, remove supernatant

4) Wash beads in 123 µl of fresh 70% EtOH twice out of magnet
5) Magnet separation, remove supernatant
6) Let air-dry at RT for 3-10 minutes (until beads stop shining, but do NOT over-dry them!)
7) Elute with 27 µL RNase/DNase free water
8) Take 1µl to run BioAnalyzer to determine how many PCR cycles for next step.

**12. PCR Enrichment**

- Use 6-9 cycles TOTAL if use started from 10-100 ng of poly-A RNA (NO depletion protocol, no pooling). Use 11-14 total cycles if you used 50-100 ng of RNA and depleted rRNAs or used 0.1-5 ng of poly-A RNA or RNA w/o depletion.
- Include a negative control for each primer set

1) Make a mix consisting of:
(P5 Enrichment Primer (universal), P7 Enrichment Primer (barcoded), see Appendix for sequence information)

| **PCR Mix** | **1 reaction** |
|---|---|
| cDNA+ (from above) | 23 µL |
| Primer mix, 25µM each (P5 Enrichment Primer + P7 Enrichment Primer) | 2 µL |
| NEBNext Q5 HotStart mix | 25 µL |
| **Total** | 50µL |

2) Run PCR with the following conditions:

| | | |
|---|---|---|
| 98°C | 40 Sec | |
| 98°C | 20 sec | 6 cycles |
| 69°C | 30 sec | |
| 72°C | 30sec | |
| 98°C | 12 sec | 9 cycles |
| 72°C | 1 min | |
| 72°C | 5 min | |

**13. SPRI Library cleanup:**

– Use 50 µl of SPRI beads (XP)/sample. Use 40-45 µl of SPRI beads if RNA fragments were LONG. Use 50-80 µl of SPRI beads/sample if RNA fragments were SHORT/extra short.

1) Add SPRI beads, mix well by slowly pipetting 10-30 times
2) Incubate at RT for 3-10 minutes
3) Place on magnet for 3-5 minutes or until solution is clear
4) Remove and discard the clear supernatant
5) Add 200 µL fresh 70-75% EtOH without removing from magnet and incubate for 30 seconds.
6) Remove and discard the clear supernatant
7) Add 200 µL fresh 70% EtOH without removing from magnet and incubate for 30 seconds

8) Remove and discard the clear supernatant
9) Air dry for 2-5 minutes
10) Elute with 21 µl of $H_2O$
11) Take 1 µl to check on BioAnalyzer

### APPENDIX: Sequences of Oligos Used in this Protocol

#### Barcoded Oligos: (regular desalting is sufficient)

- The barcoded RNA oligos and 3Tr3 second adapter oligo require a 5' phosphate and 3' blocking group (either a 3'-spacer C3 or ddC)
  e.g., 3Tr3: 5'-/5Phos/AGA TCG GAA GAG CAC ACG TCT G/3SpC3/-3'
- No special modifications for the AR2 RT primer oligo (regular desalting):
  5'-TAC ACG ACG CTC TTC CGA T-3'

#### Set of 54 RNA barcoded oligos:

| Barcode | Sequence | 6 base Barcode Read + T |
|---|---|---|
| rUrU rGrCrU rU | /5Phos/rArUrU rGrCrU rUrArG rArUrC rGrGrA rArGrA rGrCrG rUrCrG rUrGrU rArG/3SpC3/ | AAGCAAT |
| rUrG rArArU rU | /5Phos/rArUrG rArArU rUrArG rArUrC rGrGrA rArGrA rGrCrG rUrCrG rUrGrU rArG/3SpC3/ | AATTCAT |
| rArC rUrUrG rU | /5Phos/rArArC rUrUrG rUrArG rArUrC rGrGrA rArGrA rGrCrG rUrCrG rUrGrU rArG/3SpC3/ | ACAAGTT |
| rGrG rCrUrG rU | /5Phos/rArGrG rCrUrG rUrArG rArUrC rGrGrA rArGrA rGrCrG rUrCrG rUrGrU rArG/3SpC3/ | ACAGCCT |
| rUrC rArGrG rU | /5Phos/rArUrC rArGrG rUrArG rArUrC rGrGrA rArGrA rGrCrG rUrCrG rUrGrU rArG/3SpC3/ | ACCTGAT |
| rArU rUrArG rU | /5Phos/rArArU rUrArG rUrArG rArUrC rGrGrA rArGrA rGrCrG rUrCrG rUrGrU rArG/3SpC3/ | ACTAATT |
| rUrU rGrArG rU | /5Phos/rArUrU rGrArG rUrArG rArUrC rGrGrA rArGrA rGrCrG rUrCrG rUrGrU rArG/3SpC3/ | ACTCAAT |
| rUrG rGrUrC rU | /5Phos/rArUrG rGrUrC rUrArG rArUrC rGrGrA rArGrA rGrCrG rUrCrG rUrGrU rArG/3SpC3/ | AGACCAT |
| rGrU rUrUrA rU | /5Phos/rArGrU rUrUrA rUrArG rArUrC rGrGrA rArGrA rGrCrG rUrCrG rUrGrU rArG/3SpC3/ | ATAAACT |
| rArU rGrUrA rU | /5Phos/rArArU rGrUrA rUrArG rArUrC rGrGrA rArGrA rGrCrG rUrCrG rUrGrU rArG/3SpC3/ | ATACATT |
| rArU rUrGrA rU | /5Phos/rArArU rUrGrA rUrArG rArUrC rGrGrA rArGrA rGrCrG rUrCrG rUrGrU rArG/3SpC3/ | ATCAATT |
| rGrU rGrGrA rU | /5Phos/rArGrU rGrGrA rUrArG rArUrC rGrGrA rArGrA rGrCrG rUrCrG rUrGrU rArG/3SpC3/ | ATCCACT |
| rGrA rArGrA rU | /5Phos/rArGrA rArGrA rUrArG rArUrC rGrGrA rArGrA rGrCrG rUrCrG rUrGrU rArG/3SpC3/ | ATCTTCT |
| rGrG rUrCrA rU | /5Phos/rArGrG rUrCrA rUrArG rArUrC rGrGrA rArGrA rGrCrG rUrCrG rUrGrU rArG/3SpC3/ | ATGACCT |
| rGrA rGrCrA rU | /5Phos/rArGrA rGrCrA rUrArG rArUrC rGrGrA rArGrA rGrCrG rUrCrG rUrGrU rArG/3SpC3/ | ATGCTCT |
| rCrA rGrUrU rG | /5Phos/rArCrA rGrUrU rGrArG rArUrC rGrGrA rArGrA rGrCrG rUrCrG rUrGrU rArG/3SpC3/ | CAACTGT |
| rGrG rUrGrU rG | /5Phos/rArGrG rUrGrU rGrArG rArUrC rGrGrA rArGrA rGrCrG rUrCrG rUrGrU rArG/3SpC3/ | CACACCT |
| rGrA rGrGrU rG | /5Phos/rArGrA rGrGrU rGrArG rArUrC rGrGrA rArGrA rGrCrG rUrCrG rUrGrU rArG/3SpC3/ | CACCTCT |
| rUrU rCrGrU rG | /5Phos/rArUrU rCrGrU rGrArG rArUrC rGrGrA rArGrA rGrCrG rUrCrG rUrGrU rArG/3SpC3/ | CACGAAT |
| rUrC rArGrU rG | /5Phos/rArUrC rArGrU rGrArG rArUrC rGrGrA rArGrA rGrCrG rUrCrG rUrGrU rArG/3SpC3/ | CACTGAT |
| rArA rGrCrU rG | /5Phos/rArArA rGrCrU rGrArG rArUrC rGrGrA rArGrA rGrCrG rUrCrG rUrGrU rArG/3SpC3/ | CAGCTTT |
| rGrA rCrUrG rG | /5Phos/rArGrA rCrUrG rGrArG rArUrC rGrGrA rArGrA rGrCrG rUrCrG rUrGrU rArG/3SpC3/ | CCAGTCT |
| rArU rCrGrG rG | /5Phos/rArArU rCrGrG rGrArG rArUrC rGrGrA rArGrA rGrCrG rUrCrG rUrGrU rArG/3SpC3/ | CCCGATT |
| rArG rUrCrG rG | /5Phos/rArArG rUrCrG rGrArG rArUrC rGrGrA rArGrA rGrCrG rUrCrG rUrGrU rArG/3SpC3/ | CCGACTT |

| | | |
|---|---|---|
| rUrA rUrArG rG | /5Phos/rArUrA rUrArG rGrArG rArUrC rGrGrA rArGrA rGrCrG rUrCrG rUrGrU rArG/3SpC3/ | CCTATAT |
| rCrG rGrArG rG | /5Phos/rArCrG rGrArG rGrArG rArUrC rGrGrA rArGrA rGrCrG rUrCrG rUrGrU rArG/3SpC3/ | CCTCCGT |
| rCrU rArArG rG | /5Phos/rArCrU rArArG rGrArG rArUrC rGrGrA rArGrA rGrCrG rUrCrG rUrGrU rArG/3SpC3/ | CCTTAGT |
| rGrU rGrGrU rC | /5Phos/rArGrU rGrGrU rCrArG rArUrC rGrGrA rArGrA rGrCrG rUrCrG rUrGrU rArG/3SpC3/ | GACCACT |
| rGrG rGrArU rC | /5Phos/rArGrG rGrArU rCrArG rArUrC rGrGrA rArGrA rGrCrG rUrCrG rUrGrU rArG/3SpC3/ | GATCCCT |
| rGrA rGrUrG rC | /5Phos/rArGrA rGrUrG rCrArG rArUrC rGrGrA rArGrA rGrCrG rUrCrG rUrGrU rArG/3SpC3/ | GCACTCT |
| rArU rGrGrG rC | /5Phos/rArArU rGrGrG rCrArG rArUrC rGrGrA rArGrA rGrCrG rUrCrG rUrGrU rArG/3SpC3/ | GCCCATT |
| rGrU rArUrC rC | /5Phos/rArGrU rArUrC rCrArG rArUrC rGrGrA rArGrA rGrCrG rUrCrG rUrGrU rArG/3SpC3/ | GGATACT |
| rGrU rUrGrC rC | /5Phos/rArGrU rUrGrC rCrArG rArUrC rGrGrA rArGrA rGrCrG rUrCrG rUrGrU rArG/3SpC3/ | GGCAACT |
| rGrA rGrGrC rC | /5Phos/rArGrA rGrGrC rCrArG rArUrC rGrGrA rArGrA rGrCrG rUrCrG rUrGrU rArG/3SpC3/ | GGCCTCT |
| rCrA rArGrC rC | /5Phos/rArCrA rArGrC rCrArG rArUrC rGrGrA rArGrA rGrCrG rUrCrG rUrGrU rArG/3SpC3/ | GGCTTGT |
| rGrG rGrUrA rC | /5Phos/rArGrG rGrUrA rCrArG rArUrC rGrGrA rArGrA rGrCrG rUrCrG rUrGrU rArG/3SpC3/ | GTACCCT |
| rGrC rArArA rC | /5Phos/rArGrC rArArA rCrArG rArUrC rGrGrA rArGrA rGrCrG rUrCrG rUrGrU rArG/3SpC3/ | GTTTGCT |
| rUrG rArUrU rA | /5Phos/rArUrG rArUrU rArArG rArUrC rGrGrA rArGrA rGrCrG rUrCrG rUrGrU rArG/3SpC3/ | TAATCAT |
| rArC rUrGrU rA | /5Phos/rArArC rUrGrU rArArG rArUrC rGrGrA rArGrA rGrCrG rUrCrG rUrGrU rArG/3SpC3/ | TACAGTT |
| rUrA rUrGrU rA | /5Phos/rArUrA rUrGrU rArArG rArUrC rGrGrA rArGrA rGrCrG rUrCrG rUrGrU rArG/3SpC3/ | TACATAT |
| rCrA rGrGrU rA | /5Phos/rArCrA rGrGrU rArArG rArUrC rGrGrA rArGrA rGrCrG rUrCrG rUrGrU rArG/3SpC3/ | TACCTGT |
| rGrU rCrGrU rA | /5Phos/rArGrU rCrGrU rArArG rArUrC rGrGrA rArGrA rGrCrG rUrCrG rUrGrU rArG/3SpC3/ | TACGACT |
| rArG rUrArU rA | /5Phos/rArArG rUrArU rArArG rArUrC rGrGrA rArGrA rGrCrG rUrCrG rUrGrU rArG/3SpC3/ | TATACTT |
| rGrU rGrArU rA | /5Phos/rArGrU rGrArU rArArG rArUrC rGrGrA rArGrA rGrCrG rUrCrG rUrGrU rArG/3SpC3/ | TATCACT |
| rUrU rUrUrG rA | /5Phos/rArUrU rUrUrG rArArG rArUrC rGrGrA rArGrA rGrCrG rUrCrG rUrGrU rArG/3SpC3/ | TCAAAAT |
| rGrA rUrGrG rA | /5Phos/rArGrA rUrGrG rArArG rArUrC rGrGrA rArGrA rGrCrG rUrCrG rUrGrU rArG/3SpC3/ | TCCATCT |
| rArC rArGrG rA | /5Phos/rArArC rArGrG rArArG rArUrC rGrGrA rArGrA rGrCrG rUrCrG rUrGrU rArG/3SpC3/ | TCCTGTT |
| rGrG rUrUrC rA | /5Phos/rArGrG rUrUrC rArArG rArUrC rGrGrA rArGrA rGrCrG rUrCrG rUrGrU rArG/3SpC3/ | TGAACCT |
| rArU rGrUrC rA | /5Phos/rArArU rGrUrC rArArG rArUrC rGrGrA rArGrA rGrCrG rUrCrG rUrGrU rArG/3SpC3/ | TGACATT |
| rArU rArGrC rA | /5Phos/rArArU rArGrC rArArG rArUrC rGrGrA rArGrA rGrCrG rUrCrG rUrGrU rArG/3SpC3/ | TGCTATT |
| rGrG rArCrC rA | /5Phos/rArGrG rArCrC rArArG rArUrC rGrGrA rArGrA rGrCrG rUrCrG rUrGrU rArG/3SpC3/ | TGGTCCT |
| rGrU rArArC rA | /5Phos/rArGrU rArArC rArArG rArUrC rGrGrA rArGrA rGrCrG rUrCrG rUrGrU rArG/3SpC3/ | TGTTACT |
| rCrG rGrGrA rA | /5Phos/rArCrG rGrGrA rArArG rArUrC rGrGrA rArGrA rGrCrG rUrCrG rUrGrU rArG/3SpC3/ | TTCCCGT |
| rGrC rGrGrA rA | /5Phos/rArGrC rGrGrA rArArG rArUrC rGrGrA rArGrA rGrCrG rUrCrG rUrGrU rArG/3SpC3/ | TTCCGCT |

#### Library Enrichment Primers (regular desalting):

##### P5 Enrichment Primer (universal):

5'–AATGATACGGCGACCACCGAGATCTACACTCTTTCCCTACACGACGCTCTTCCGATCT-3'

##### P7 Enrichment Primer (barcoded):

| P7 Enrichment Primer Sequence (5' --> 3') with barcode | barcodes (BC) | BC READ (reverse complement) |
|---|---|---|
| CAAGCAGAAGACGGCATACGAGATTCGTGTGCGTGACTGGAGTTCAGACGTGTGCTCTTCCGATCT | **TCGTGTGC** | **GCACACGA** |
| CAAGCAGAAGACGGCATACGAGATTCGCCAGAGTGACTGGAGTTCAGACGTGTGCTCTTCCGATCT | **TCGCCAGA** | **TCTGGCGA** |
| CAAGCAGAAGACGGCATACGAGATTCGCTATGGTGACTGGAGTTCAGACGTGTGCTCTTCCGATCT | **TCGCTATG** | **CATAGCGA** |
| CAAGCAGAAGACGGCATACGAGATGGCTCCTGGTGACTGGAGTTCAGACGTGTGCTCTTCCGATCT | **GGCTCCTG** | **CAGGAGCC** |
| CAAGCAGAAGACGGCATACGAGATATCCGACAGTGACTGGAGTTCAGACGTGTGCTCTTCCGATCT | **ATCCGACA** | **TGTCGGAT** |
| CAAGCAGAAGACGGCATACGAGATAACATAATGTGACTGGAGTTCAGACGTGTGCTCTTCCGATCT | AACATAAT | **ATTATGTT** |
| CAAGCAGAAGACGGCATACGAGATATGGTAGGGTGACTGGAGTTCAGACGTGTGCTCTTCCGATCT | ATGGTAGG | **CCTACCAT** |
| CAAGCAGAAGACGGCATACGAGATGCTAAGTAGTGACTGGAGTTCAGACGTGTGCTCTTCCGATCT | GCTAAGTA | **TACTTAGC** |
